# CHC22 clathrin mediates traffic from early secretory compartments for human GLUT4 pathway biogenesis

**DOI:** 10.1101/242941

**Authors:** Stéphane M. Camus, Marine D. Camus, Carmen Figueras-Novoa, Gaelle Boncompain, L. Amanda Sadacca, Christopher Esk, Anne Bigot, Gwyn W. Gould, Dimitrios Kioumourtzoglou, Franck Perez, Nia J. Bryant, Shaeri Mukherjee, Frances M. Brodsky

## Abstract

Glucose Transporter 4 (GLUT4) is sequestered inside muscle and fat, then released by vesicle traffic to the cell surface in response to post-prandial insulin for blood glucose clearance. Here we map the biogenesis of this GLUT4 traffic pathway in humans, which involves clathrin isoform CHC22. We observe that GLUT4 transits through the early secretory pathway more slowly than the constitutively-secreted GLUT1 transporter and localize CHC22 to the endoplasmic-reticulum-to-Golgi-intermediate compartment (ERGIC). CHC22 functions in transport from the ERGIC, as demonstrated by an essential role in forming the replication vacuole of *Legionella pneumophila* bacteria, which requires ERGIC-derived membrane. CHC22 complexes with ERGIC tether p115, GLUT4 and sortilin and down-regulation of either p115 or CHC22, but not GM130 or sortilin abrogate insulin-responsive GLUT4 release. This indicates CHC22 traffic initiates human GLUT4 sequestration from the ERGIC, and defines a role for CHC22 in addition to retrograde sorting of GLUT4 after endocytic recapture, enhancing pathways for GLUT4 sequestration in humans relative to mice, which lack CHC22.

**Summary:** Blood glucose clearance relies on insulin-mediated exocytosis of glucose transporter 4 (GLUT4) from sites of intracellular sequestration. We show that in humans, CHC22 clathrin mediates membrane traffic from the ER-to-Golgi Intermediate Compartment, which is needed for GLUT4 sequestration during GLUT4 pathway biogenesis.

---

Glucose transporter 4 (GLUT4) mediates post-prandial blood glucose clearance into muscle and adipose tissues following insulin-stimulated translocation to the cell surface from sites of intracellular sequestration, known collectively as the GLUT4 storage compartment (GSC) (Bogan, 2012; Leto and Saltiel, 2012). Deregulation of GLUT4 vesicle release occurs during insulin resistance and contributes to pathogenesis of type 2 diabetes (T2D) (Bogan, 2012). In rodent models, endocytic pathways have been identified as essential routes for retrograde recycling of GLUT4 to reform insulin-responsive vesicles after insulin-mediated release (Antonescu et al., 2008; Bryant et al., 2002; Fazakerley et al., 2009; Jaldin-Fincati et al., 2017; Kandror and Pilch, 2011). Endosomal sorting and retrograde transport through the trans-Golgi network is involved in this process, generating the GSC (Shewan et al., 2003), which is a mixture of tubules and vesicles in which GLUT4 is sequestered in the absence of insulin. The trafficking routes by which newly synthesized GLUT4 accesses the GSC and participates in its formation are less well defined. In human myocytes and adipocytes, GSC formation involves the non-canonical isoform of clathrin, CHC22, which is missing from rodents due to loss of the encoding gene (Wakeham et al., 2005). Here we define a role for CHC22 clathrin in the biosynthetic trafficking pathway delivering GLUT4 to the GSC in humans.

The non-canonical clathrin isoform CHC22 is encoded on human chromosome 22 and has 85% sequence identity with the canonical CHC17 clathrin isoform (Wakeham et al., 2005). CHC17 performs receptor-mediated endocytosis at the plasma membrane and protein sorting at the trans-Golgi network in all eukaryotic cells and tissues (Brodsky, 2012). CHC22 has been implicated in distinct tissue-specific membrane traffic pathways consistent with its different biochemical properties and restricted tissue expression. While both CHC22 and CHC17 homotrimerize into triskelia that assemble to form latticed vesicle coats, the CHC22 coat is more stable and within cells, the two clathrins form separate vesicles (Dannhauser et al., 2017). CHC22 does not bind the clathrin light chain subunits associated with CHC17 or the endocytic AP2 adaptor that recruits CHC17 to the plasma membrane, while CHC22 interacts preferentially with the GGA2 adaptor compared to CHC17 (Dannhauser et al., 2017; Liu et al., 2001; Vassilopoulos et al., 2009). In agreement with its adaptor specificity, several analyses have now confirmed that CHC22 does not support receptor-mediated endocytosis at the plasma membrane (Dannhauser et al., 2017), though earlier studies suggested that it might replace CHC17 function upon over-expression (Hood and Royle, 2009).

In humans, CHC22 is expressed most highly in muscle, reaching about 10% of CHC17 levels, and has variable but lower expression in other tissues (Esk et al., 2010). In both human myocytes and adipocytes, CHC22 is needed for formation of the GSC, a membrane traffic pathway that these cell types uniquely share (Vassilopoulos et al., 2009). We previously observed that CHC22 is required for a retrograde transport pathway from endosomes (Esk et al., 2010), a step that CHC17 can also perform (Johannes and Popoff, 2008) and which has been shown to be important in murine GSC formation (Jaldin-Fincati et al., 2017). However, when CHC22 is depleted from human myocytes, CHC17 does not compensate for CHC22 loss and cells are unable to form an insulin-responsive GSC, suggesting that CHC22 mediates an additional pathway in human GSC formation (Vassilopoulos et al., 2009). CHC22 is also transiently expressed in the developing human brain (Nahorski et al., 2015) and has been implicated in protein targeting to dense core secretory granules, another pathway that involves sequestration of cargo away from standard endocytic and secretory pathways (Nahorski et al., 2018).

In the adipocytes and myocytes of insulin resistant type-2 diabetic patients, GLUT4 accumulates intracellularly (Garvey et al., 1998; Maianu et al., 2001), in a region where CHC22 also accumulates (Vassilopoulos et al., 2009). Transgenic expression of CHC22 in murine muscle caused similar accumulation of GLUT4 with CHC22 along with two other proteins involved in intracellular GLUT4 sorting, IRAP and VAMP2, and aged CHC22-transgenic animals developed hyperglycemia. These observations not only highlight fundamental differences in GLUT4 intracellular trafficking to the GSC between human and mice, but also link abnormal CHC22 intracellular localization and function to defects in GLUT4 trafficking during insulin resistance. Therefore, mapping the CHC22-mediated GLUT4 trafficking pathways leading to the biogenesis of the GSC in humans is relevant to pathophysiology leading to type 2 diabetes. Understanding CHC22’s role in GLUT4 traffic should also shed light on its role during the development of pain-sensing neurons, which was found to be defective in children homozygous for a rare familial mutation in the CHC22-encoding gene, who unfortunately did not survive to an age where their glucose metabolism could be studied (Nahorski et al., 2015).

In the present study, we identify a specialized pathway that CHC22 mediates during biogenesis of the human GSC by analyzing CHC22 function and distribution in several human cell models. We observed that CHC22 localizes to the early part of the secretory pathway where GLUT4 is delayed during its biogenesis relative to the constitutively-secreted GLUT1 transporter. In particular, CHC22 co-localizes and complexes with p115, a resident of the ER-to-Golgi Intermediate Compartment (ERGIC) where it participates in membrane export from that compartment. We confirmed that this compartment was the ERGIC, by utilizing the bacterium *Legionella pneumophila*, which is known to co-opt membrane from the ERGIC to avoid the degradative environment of the endocytic pathway. Along with CHC22, the bacterial compartment also acquired essential components of the GLUT4 pathway, namely IRAP, sortilin and GGA2. We further found that the CHC22-dependent pathway for GLUT4 transport to the human GSC relies on p115 but not GM130, indicating that this pathway depends on membrane traffic from the ERGIC in an unconventional secretory route for intracellular sequestration.

## Results

### HeLa-GLUT4 cells have an insulin-responsive GLUT4 trafficking pathway that requires CHC22

To study the role of CHC22 in formation of the human GSC, we established a cellular model in which GLUT4 translocation can be easily detected. This was necessitated by the limited experimental capacity of available human muscle and adipocyte cell lines and the lack of reagents to detect endogenous GLUT4 at the cell surface. We generated a stable HeLa cell line (HeLa-GLUT4) expressing human GLUT4 containing a haemagglutinin (HA) tag in the first exofacial loop and a GFP tag at the intracellular carboxyl terminus, a construct that has been extensively characterized and validated for the study of intracellular GLUT4 trafficking in rodent cells (Dawson et al., 2001; Dobson et al., 1996). HeLa cells were chosen because they have levels of CHC22 comparable to those in human muscle cells (Esk et al., 2010) and they also express sortilin, another protein important for GLUT4 traffic to an insulin-responsive compartment (Huang et al., 2013; Shi and Kandror, 2005; Shi and Kandror, 2007).

GLUT4 was sequestered intracellularly in HeLa-GLUT4 cells in the absence of insulin (basal), localizing to peripheral vesicles (arrowheads) and a perinuclear depot (arrows), as observed for insulin-releasable GLUT4 vesicles and the tubulo-vesicular GSC of murine cells (Bryant et al., 2002; Shewan et al., 2003) (Fig. 1 A). Upon insulin treatment (15 min), GLUT4 was detected at the cell surface using a monoclonal antibody to the haemagglutinin (HA) tag (anti-HA) (Fig. 1 A). The degree of GLUT4 translocation was quantified using a fluorescence-activated cell sorter (FACS) to measure the mean fluorescence intensity (MFI) of surface GLUT4 relative to the MFI of total cellular GLUT4 (GFP signal) (Fig. 1 B). Additionally, treatment of HeLa-GLUT4 cells with insulin induced phosphorylation of AKT and its substrate AS160, two modifications required for insulin-stimulated GLUT4 translocation (Bogan, 2012) (Fig. 1 B). Intracellular sequestration and insulin responsiveness of GLUT4 was observed to be specific by comparison to endogenous Class I molecules of the Major Histocompatibility Complex (MHCI), which were constitutively expressed at the plasma membrane of HeLa-GLUT4 cells, clearly segregated from GLUT4 under basal conditions (Fig. 1 C), and unchanged by insulin treatment. Upon insulin treatment, GLUT4 co-localized with MHCI at the cell surface and this was prevented by treatment of HeLa-GLUT4 cells with siRNA targeting CHC22 (Fig. 1 C and D). Down-regulation of CHC22 did not affect the distribution of MHCI in either basal or insulin-stimulated conditions (Fig. 1 C). Thus formation of the insulin-responsive GLUT4 pathway in HeLa-GLUT4 cells depends on CHC22.

**Figure 1:**
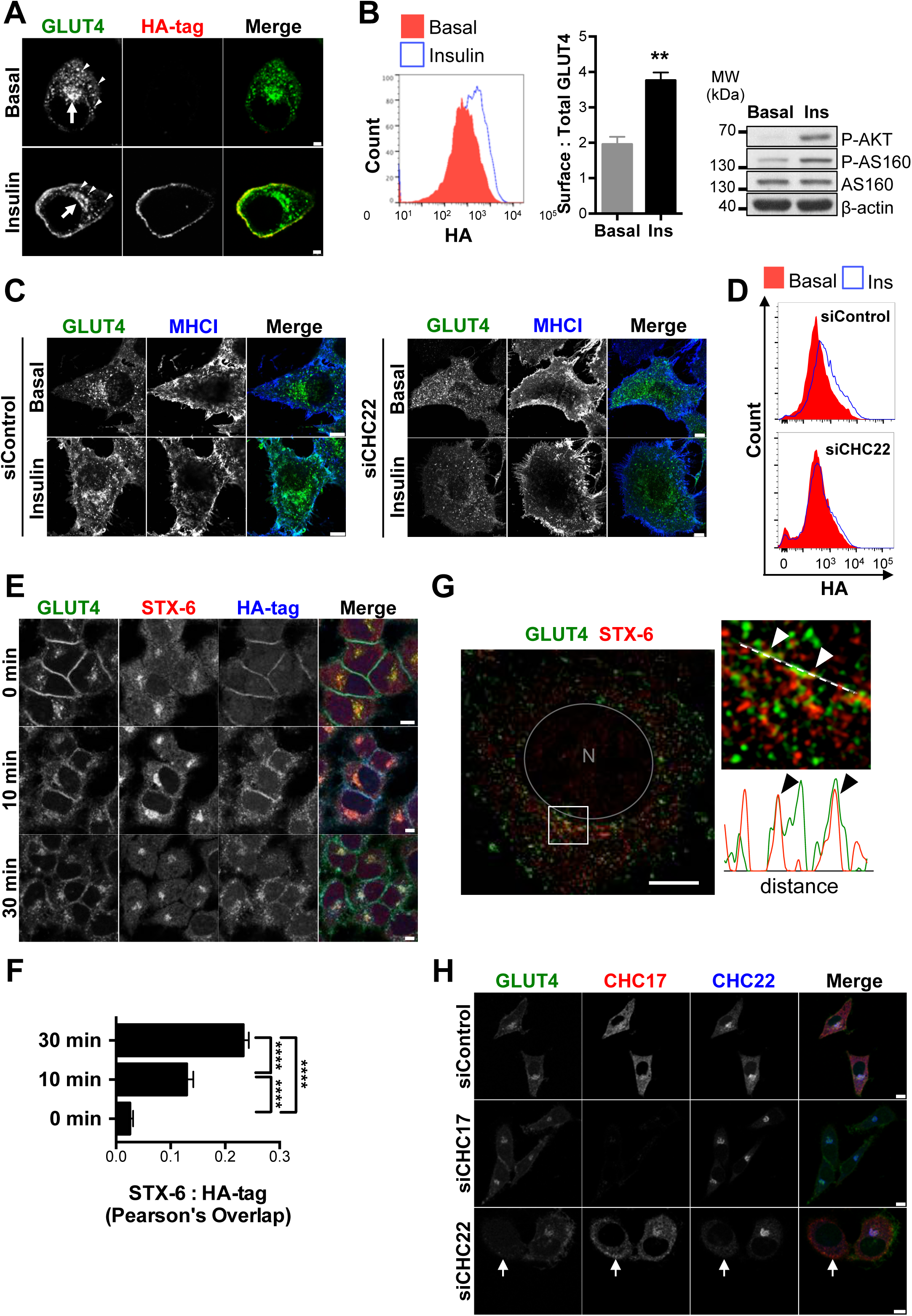
HeLa-GLUT4 cells have a functional GLUT4 trafficking pathway that requires CHC22. (A) Representative images of GLUT4 (exofacial haemagglutinin (HA)-tag, internal GFP tag) in HeLa-GLUT4 cells before (basal) or after insulin treatment. GLUT4 at the plasma membrane was detected by immunofluorescence (IF) after surface labeling with anti-HA monoclonal antibody (red). Total GLUT4 (green) was detected by GFP tag. Arrows show the GLUT4 storage compartment. Arrowheads point to peripheral GLUT4 vesicles. Scale bars: 7.5 µm. (B) Left panel – Representative FACS histogram of surface GLUT4 fluorescence intensities (signal from anti-HA labeling) before (basal) and after insulin treatment (Ins). Middle panel – Quantification of surface:total GLUT4 (HA:GFP mean fluorescence intensity signals). Data expressed as mean ± SEM, N=3, 10,000 cells acquired per experiment. Two-tailed unpaired Student’s t-test with equal variances, **p<0.01. Right panel – Representative immunoblot for phosphorylated AKT (p-AKT), phosphorylated AS160 (p-AS160), total AS160 and β-actin in HeLa-GLUT4 cells before and after insulin treatment. The migration position of molecular weight (MW) markers is indicated at the left in kilodaltons (kDa). (C) Representative images of total GLUT4 (GFP tag, green) and Major Histocompatibility Complex I molecules (MHCI, blue) before (basal) or after insulin treatment in HeLa-GLUT4 cells transfected with non-targeting control siRNA (siControl) or siRNA targeting CHC22 (siCHC22). Scale bars: 8 µm. (D) Representative FACS histograms of surface GLUT4 fluorescence intensity (signal from anti-HA labeling) in HeLa-GLUT4 cells transfected with siControl or siRNA siCHC22 before (red) or after treatment with insulin (blue). Histograms are extracted from the experiment quantified in Fig. 8 A. (E) Representative IF staining for internalized surface-labeled GLUT4 (HA-tag, blue) and syntaxin 6 (STX-6, red) for HeLa-GLUT4 cells at 0, 10 or 30 minutes after insulin treatment. Total GLUT4 is detected by GFP tag (green). Scale bars: 7.5 µm. (F) Pearson’s overlap quantification for labeling of STX-6 and HA-tag (Data expressed as mean ± SEM, N=3, 14-19 cells per experiment). One-way analysis of variance (ANOVA) followed by Bonferroni’s multiple comparison post-hoc test ****p<0.0001. (G) Left panel – representative Structured Illumination Microscopy (SIM) image of a HeLa-GLUT4 cell stained for STX-6 (red). Total GLUT4 (green) was detected by GFP tag. The gray circle delineates the nucleus (N) and the white square delineates the magnified area displayed in the right panel image. Scale bar: 10 µm. Right panel – the white dashed line in the magnified area spans the segment for which fluorescence intensities for GLUT4 and STX-6 are plotted below, in green and red, respectively. Arrowheads indicate areas of overlap. (H) Representative IF staining for CHC17 (red) and CHC22 (blue) in HeLa-GLUT4 cells transfected with non-targeting siControl or siRNA targeting CHC17 (siCHC17) or siCHC22, with GLUT4 detected by GFP tag (green). Arrows point to a CHC22-depleted cell. Scale bars: 10 µm for siControl and siCHC17 and 7.5 µm for siCHC22. Merged images in (A), (C), (E), (G), (H) show red/green overlap in yellow, red/blue overlap in magenta, green/blue overlap in turquoise, and red/green/blue overlap in white.

In rodent cells, GLUT4 released by insulin to the plasma membrane can return by endocytosis and retrograde transport to a syntaxin 6 (STX-6)–positive GSC within 30 minutes (Perera et al., 2003). Replicating this pulse-chase experiment in the HeLa-GLUT4 model, we labeled surface GLUT4 with anti-HA antibody following insulin stimulation and tracked internalized GLUT4 to a perinuclear compartment overlapping with STX-6, with similar kinetics to those observed in rodent cells (Fig. 1 E and F). Using structured-illumination microscopy (SIM), we observed that, under basal conditions, the perinuclear GLUT4 depot in HeLa-GLUT4 cells partially co-localized with STX-6 (arrowheads, Fig. 1 G), which is considered a marker of the GSC in rodent cells (3T3-L1 mouse adipocytes and L6 rat myotubes) (Foley and Klip, 2014; Shewan et al., 2003). The GSC in HeLa-GLUT4 cells also had properties of the human GSC in that siRNA-mediated depletion of CHC22 induced dispersal of GLUT4 from the perinuclear region (Fig. 1 H) and inhibited insulin-stimulated GLUT4 translocation (Fig. 1 D)(Esk et al., 2010; Vassilopoulos et al., 2009). Taken together, our results show that HeLa cells stably expressing HA-GLUT4-GFP form an intracellular GSC, respond to insulin stimulation by rapidly translocating GLUT4 to the plasma membrane and recycle GLUT4 back to the GSC. Furthermore, CHC22 expression is critical for establishing this GLUT4 trafficking pathway. Thus, the model recapitulates features of the GLUT4 pathway observed for both mouse and human cells and, while the pathway may not have every regulator controlling GLUT4 traffic in muscle and adipocytes, it can be studied to identify players and routes involved in GLUT4 trafficking in human cells. We note that during the course of this work, other laboratories developed and validated similar models of insulin-dependent GLUT4 translocation in HeLa cells (Haga et al., 2011; Trefely et al., 2015).

### Newly synthesized GLUT4 co-localizes with CHC22 and is delayed in the early secretory pathway relative to constitutively expressed GLUT1

Formation of the GSC in rodent cells relies on the biosynthetic pathway feeding the GSC with newly synthesized GLUT4 (Watson et al., 2004) and recycling pathways that replenish it after insulin-mediated GLUT4 exocytosis (Antonescu et al., 2008; Bryant et al., 2002; Fazakerley et al., 2009; Kandror and Pilch, 2011; Martin et al., 2006). Our previous studies demonstrated a role for CHC22 in human GSC formation (Vassilopoulos et al., 2009) and identified a function for CHC22 in retrograde transport from endosomes (Esk et al., 2010), suggesting that CHC22 could participate in GSC replenishment after GLUT4 translocation. To address whether CHC22 is also involved in biogenesis of the GLUT4 pathway from the secretory pathway, we tracked newly synthesized GLUT4 relative to the constitutively-expressed GLUT1, (Hresko et al., 1994; Hudson et al., 1992). Their biosynthetic pathways were compared by the Retention Using Selective Hooks (RUSH) approach (Boncompain et al., 2012) for which each transporter was tagged with a streptavidin-binding protein (SBP) as well as GFP or mCherry and an HA tag. The resulting fusion proteins HA-GLUT4-SBP-GFP and HA-GLUT1-SBP-mCherry, were co-expressed in HeLa cells also expressing streptavidin fused to an ER-resident isoform of the human invariant chain (Ii-hook) (Schutze et al., 1994). The ER-retained HA-GLUT1-SBP-mCherry and HA-GLUT4-SBP-GFP were then synchronously released for ongoing traffic upon addition of biotin to the cells (Fig. 2 A, Videos 1 and 2). Following release, both transporters showed initial perinuclear localization, but after 25 minutes the GLUT1 and GLUT4 pathways diverged and vesicles with GLUT1 were observed trafficking to the cell surface (arrowheads, 26-57 minutes post-biotin addition), while GLUT4 remained concentrated in a perinuclear region. This was consistent with the expected constitutive secretion of GLUT1 and sequestration of GLUT4 and further demonstrated the specificity of intracellular sequestration of GLUT4 in HeLa cells.

**Figure 2:**
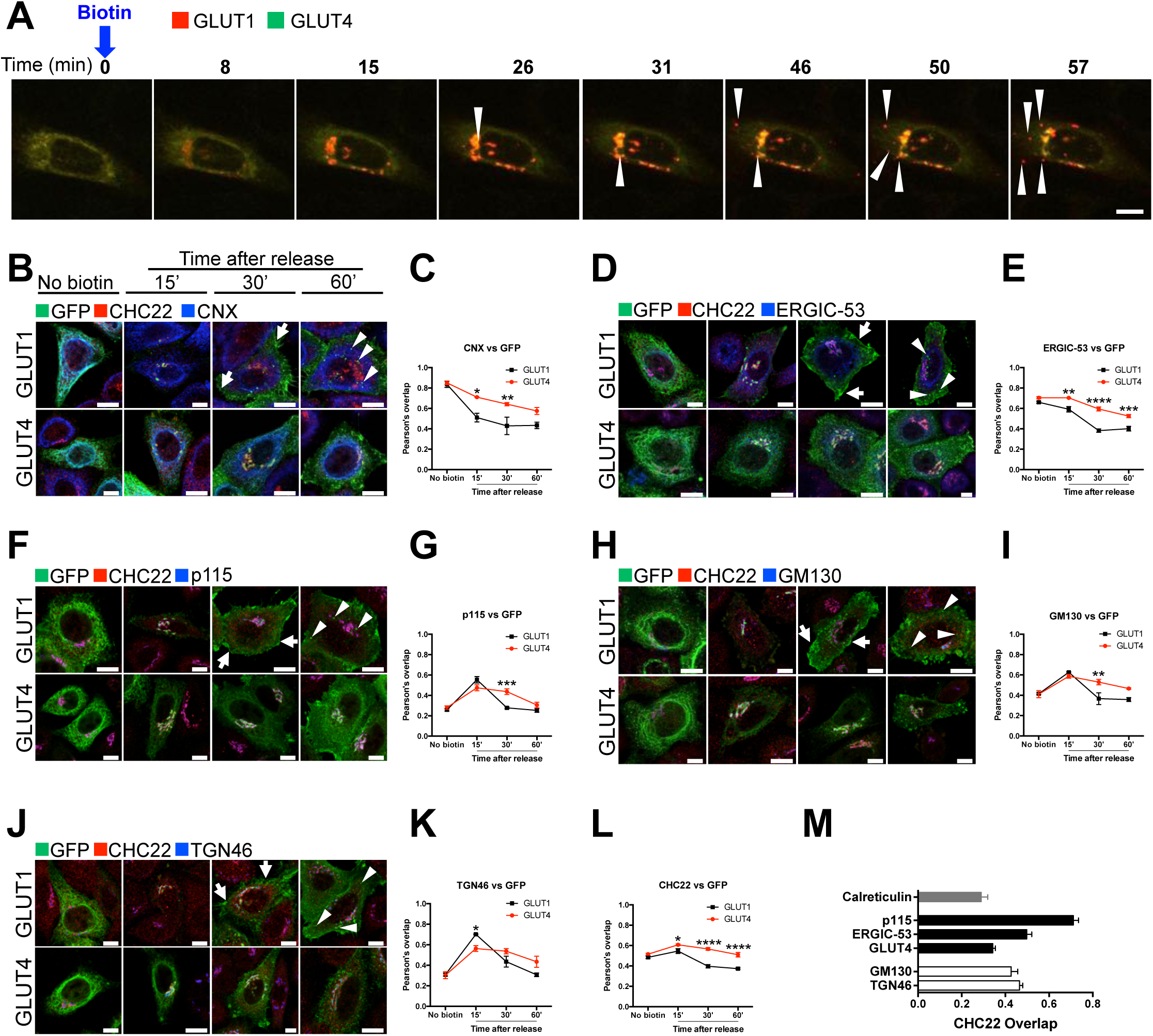
Newly synthesized GLUT4 is delayed in the early secretory pathway compared to GLUT1. (A) Representative stills extracted from Video 1 showing a HeLa cell expressing the endoplasmic reticulum (ER) Ii-hook fused to streptavidin along with HA-GLUT1-SBP-mCherry (GLUT1, red) and HA-GLUT4-SBP-GFP (GLUT4, green). The intracellular traffic of GLUT1-mCherry and GLUT4-GFP was simultaneously tracked for 1h after biotin addition released them from the ER. Upon ER exit, both GLUT1 and GLUT4 accumulated in the perinuclear region of the cell (yellow). From 26 min onwards, highly mobile GLUT1 vesicles (arrowheads) were visible (red) while GLUT4 remained perinuclear. Scale bar: 10 µm. (B, D, F, H, J) Representative immunofluorescence staining for GLUT1-SBP-GFP or GLUT4-SBP-GFP (detected with anti-GFP antibody, green), CHC22 (red) and (B) calnexin (CNX, blue), (D) ERGIC-53 (blue), (F) p115 (blue), (H) GM130 (blue) or (J) TGN46 (blue) in HeLa cells expressing HA-GLUT1-SBP-GFP or HA-GLUT4-SBP-GFP along with the ER Ii-hook. Traffic of GLUT4 and GLUT1 was tracked at 0, 15, 30 and 60 minutes after release from the ER by biotin. Arrows point to GLUT1 detected at the plasma membrane and arrowheads point to GLUT1-positive endosomal structures. Merged images show red/green overlap in yellow, red/blue overlap in magenta, green/blue overlap in turquoise, and red/green/blue overlap in white. Scale bars: 10 µm. (C, E, G, I, K, L) Pearson’s overlap between GLUT1 or GLUT4 and CNX, ERGIC-53, p115, GM130, TGN46 or CHC22 at different time-points post-ER release. Data expressed as mean ± SEM, N=3-4, 10-46 cells per experiment. One-way analysis of variance (ANOVA) followed by Sidak’s multiple comparison post-hoc test *p<0.05, **p<0.01, ****p<0.0001 to test differences between GLUT1 and GLUT4 overlap with markers at each time points. (M) Pearson’s overlap between CHC22 and GLUT4, ER marker calreticulin, ERGIC markers p115 and ERGIC-53, cis-Golgi marker GM130 or trans-Golgi marker TGN46 in HeLa-GLUT4 from images taken by confocal microscopy (corresponding representative immunofluorescence staining in Fig. S1). Data expressed as mean ± SEM, N=3, 4-10 cells across 3 independent samples.

We then analyzed these pathways in more detail by visualizing the traffic of HA-GLUT4-SBP-GFP or HA-GLUT1-SBP-GFP relative to CHC22 and to markers of the secretory pathway at different time points after biotin addition, and co-localization was quantified. Within 30 minutes of biotin addition, GLUT1 rapidly exited the ER and trafficked through the Golgi apparatus as exemplified by the 40% decrease in overlap with the ER resident protein calnexin (CNX) (Fig. 2, B and C; black line) and the transient increase in overlap with ERGIC markers (p115 and ERGIC-53, Fig. 2 D-G; black lines), the Golgi marker GM130 and the trans-Golgi marker TGN46 (Fig. 2, H-K; black lines). GLUT1 was detected at the plasma membrane as soon as 30 minutes post ER release (arrows) and by 60 minutes, GLUT1 was detected in endosomal structures (arrowheads), indicating internalization from the plasma membrane. In contrast, GLUT4 exit from the ER was slower (Fig. 2, B and C, red lines). GLUT4 overlap with ERGIC-53, p115 and GM130 and TGN46 increased at 15 minutes post release but did not decrease over time (Fig. 2, D-K, red lines). GLUT4 co-localization with CHC22 was similar to its residence with secretory pathway markers, peaking at 15 minutes but remaining more co-localized with CHC22 than GLUT1, which only transiently overlapped with CHC22 (Fig. 2 L). Overall, these experiments indicate that the trafficking kinetics of newly synthesized GLUT1 and GLUT4 are fundamentally different. Moreover, following ER release, GLUT4, and not GLUT1, was retained in a perinuclear region that overlaps with CHC22, suggesting a role for CHC22 in trafficking newly synthesized GLUT4 and that CHC22 might interact with secretory pathway compartments.

### CHC22 localizes with markers of the ER-to-Golgi Intermediate Compartment

To identify potential locations for CHC22 function in transporting newly synthesized GLUT4, we analyzed CHC22 overlap with markers of the secretory pathway in HeLa-GLUT4 cells and in human myotubes formed by differentiation of two different human myoblast cell lines (LHCNM2 or hSkMC-AB1190-GLUT4) (Figs. 2 M, S1, 3 and 4). The LHCNM2 cells express low levels of endogenous GLUT4 upon differentiation (Vassilopoulos et al., 2009) and the hSkMC-AB1190-GLUT4 cells were derived from the transformed human myoblast (line AB1190) by transfection and selected for permanent expression of HA-GLUT4-GFP. These cells were analyzed to establish whether pathways identified in the HeLa-GLUT4 cells are present in human myotubes where the GLUT4 pathway naturally operates. Using a commercially available polyclonal antibody specific for CHC22 and not reactive with CHC17 (Fig. S1 A), we observed significant co-localization of CHC22 with two markers of the ERGIC, namely p115 (Alvarez et al., 2001) and ERGIC-53 (Lahtinen et al., 1996) in both the HeLa-GLUT4 model and in human myotubes (Figs. 2 M and S1 B and C). CHC22 only partially overlapped with Golgi markers GM130 and TGN46 (Figs. 2 M and S1 D and E) and no significant co-localization was seen with ER markers calreticulin or calnexin in all cell lines (Figs. 2 M and S1 F and G). Given the limited spatial resolution of conventional laser scanning confocal microscopy (200 nm), these overlap values could be overestimated. We therefore used super-resolution Structured Illumination Microscopy (SIM), which improves lateral resolution twofold (100 nm). Using SIM, we confirmed the substantial overlap between CHC22 and ERGIC markers p115 and ERGIC-53 in HeLa-GLUT4 and in the human skeletal muscle cells (Figs. 3 A-D, arrowheads, and S1 I) with most extensive overlap between CHC22 and p115. SIM analysis did not support the apparent overlap between CHC22 and Golgi markers obtained by confocal microscopy in either HeLa-GLUT4 or differentiated human myoblasts. In both cell types, CHC22 was separated from cis-Golgi (GM130) and trans-Golgi (TGN46) markers (Figs. 4 A and B and S1 I), though localized adjacent to both compartments. Co-staining for p115 and STX-6 showed alignment of compartments with these markers in both cell types but little overlap (Fig. 4 C), suggesting that ERGIC and CHC22 compartments are closely associated with but not coincident with sites of GLUT4 retrograde transport. GLUT4 was widely distributed in the cells analyzed by SIM without preferential co-localization with any particular marker analyzed.

**Figure 3:**
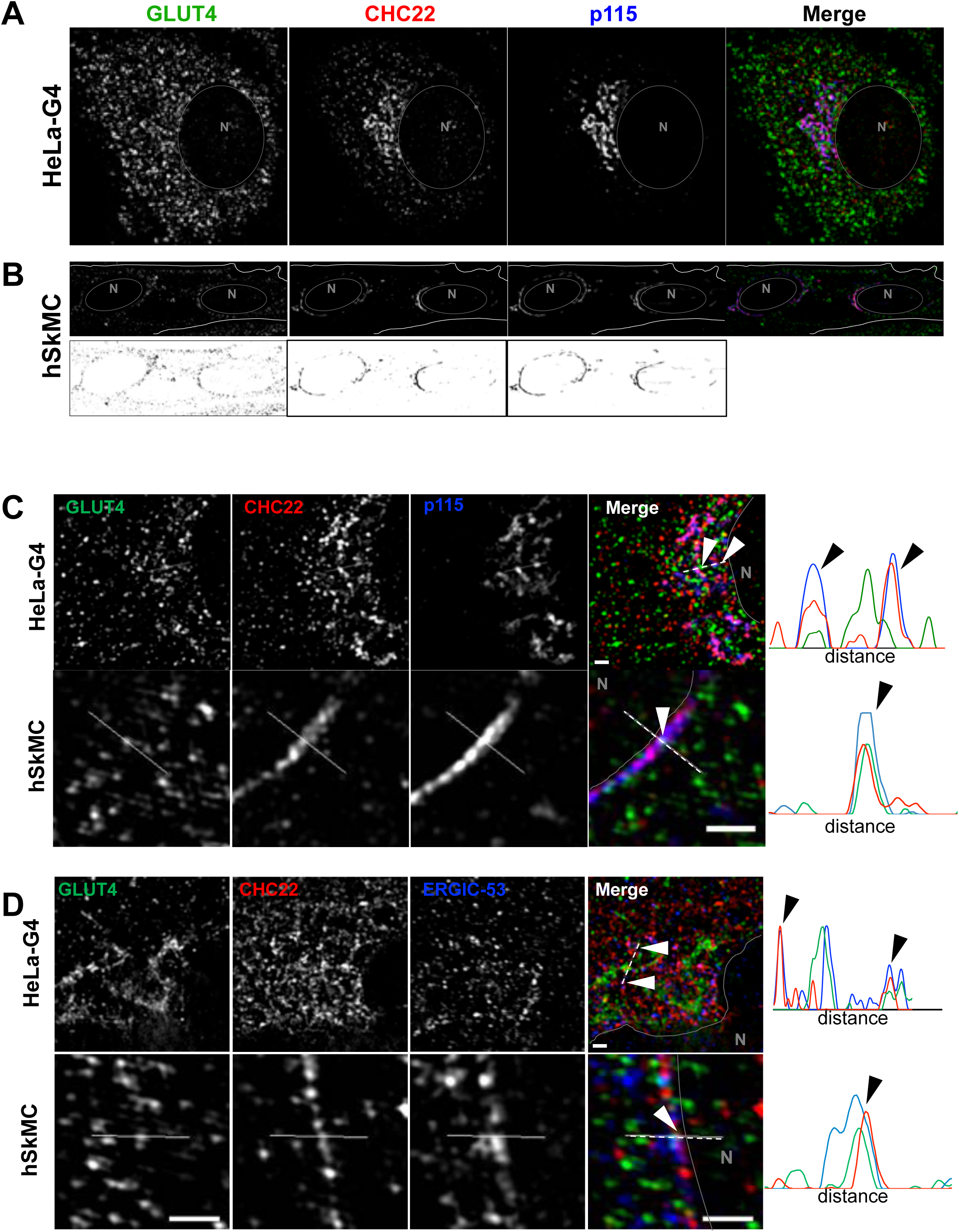
CHC22 is localized at the ER-to-Golgi Intermediate Compartment in HeLa GLUT4 and human myotubes. (A and B) Representative Structured Illumination Microscopy of a HeLa-GLUT4 cell (HeLa-G4) (A) and the human skeletal muscle cell line hSkMC-AB1190-GLUT4 (hSkMC) (B) stained for CHC22 (red) and p115 (blue). The gray circles delineate the nuclei (N). Muscle cell staining with each antibody is shown in black on white below the color images. Scale bars: 10 µm. (C and D) Representative Structured Illumination Microscopy of the perinuclear region of HeLa-GLUT4 cells and hSkMC-AB1190-GLUT4 stained for CHC22 (red) and p115 (C), ERGIC-53 (blue) (D). The solid gray lines delineate the nuclear border (N). The dashed white lines span the segment for which fluorescence intensities for GLUT4 (green), CHC22 (red) and p115, ERGIC-53 (blue) were plotted. Arrowheads indicate areas of peak overlap. Scale bars: 1 µm. In A, B, C and D, GLUT4 (green) was detected by GFP tag in HeLa-GLUT4 or immunostained with anti-GFP antibody in hSkMC-AB1190-GLUT4. Merged images in (A, B and C) show red/green overlap in yellow, red/blue overlap in magenta, green/blue overlap in turquoise, and red/green/blue overlap in white.

**Figure 4:**
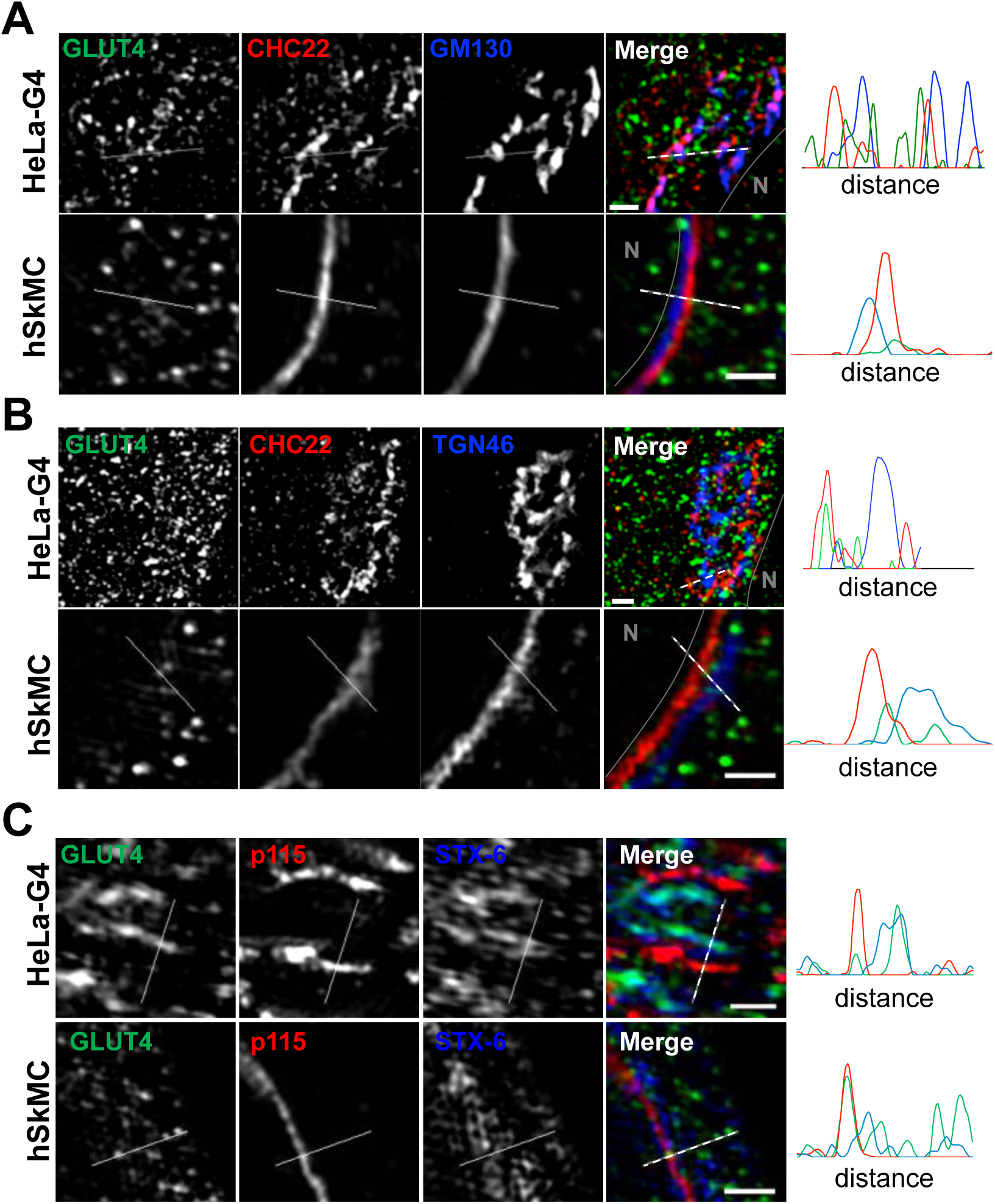
The CHC22 compartment is localized proximal to the trans-Golgi Network and does not overlap with the cis-Golgi. (A, B and C) Representative Structured Illumination Microscopy of the perinuclear region of HeLa-GLUT4 cells (HeLa-G4) and hSkMC-AB1190-GLUT4 (hSkMC) stained for CHC22 (red) and GM130 (A), TGN46 (B) and syntaxin 6 (STX-6, blue) (C). The solid gray lines delineate the nuclear border (N). The dashed white lines span the segment over which fluorescence intensities for GLUT4 (green), CHC22 (red) and GM130, TGN46 and STX-6 (blue) were plotted. Scale bars: 1 µm. In A, B and C, GLUT4 (green) was detected by GFP tag in HeLa-GLUT4 in HeLa-GLUT4 or immunostained using an anti-GFP antibody in hSkMC-AB1190-GLUT4. Merged images in (A, B and C) show red/green overlap in yellow, red/blue overlap in magenta, green/blue overlap in turquoise, and red/green/blue overlap in white.

To further define the relationship between CHC22, ERGIC markers and the retrograde transport of GLUT4 internalized after insulin-mediated release, HeLa-GLUT4 cells were treated with insulin and GLUT4 internalization was tracked by uptake of anti-HA antibody. By confocal microscopy, GLUT4 showed time-dependent co-localization with CHC22 and ERGIC-53 (Fig. S2) after 10 and 30 minutes of re-uptake, indicating that recaptured GLUT4 accumulates in close proximity to CHC22 and the ERGIC. When HeLa-GLUT4 or the hSkMC-AB1190-GLUT4 human myotubes were treated with Brefeldin A (BFA), CHC22 co-distributed with the ERGIC markers p115, ERGIC-53 and Rab1 and segregated away from the perinuclear GLUT4 (Fig. S3). This is consistent with our previous observations that BFA does not cause CHC22 to dissociate from intracellular membranes (Liu et al., 2001) and supports association of CHC22 with ERGIC membranes. We also observed that BFA treatment did not affect GLUT4 release to the plasma membrane in response to insulin and that GLUT4 was less localized with CHC22 or ERGIC markers upon insulin stimulation, in the presence or absence of BFA (Fig. S3 C-F). This supports observations from others showing that GSC formation is not affected by BFA (Martin et al., 2000). Together, these data suggest that CHC22-mediated trafficking leads to the GSC, and could be involved in initial GSC formation from the secretory pathway by mediating GLUT4 traffic emerging from the ERGIC in addition to its previously defined role in retrograde transport from endosomes to the GSC (Esk et al., 2010). Furthermore, these results highlight the close proximity of the ERGIC to the compartment where GLUT4 accumulates after endocytic recapture.

### CHC22 participates in membrane trafficking from the ERGIC

To establish whether CHC22 has functional activity at the ERGIC, we took advantage of *Legionella pneumophila* (*L.p.*), a facultative intracellular pathogen that avoids the host’s endo-lysosomal compartment and specifically hijacks membranes from the early secretory pathway to create an ER/ERGIC-like, *L.p.*-containing vacuole (LCV) for replication (Kagan and Roy, 2002). Upon infection, *L.p.* secretes ∼300 effector proteins, through a type IV secretion system, some of which enable recruitment of ER / ERGIC proteins calnexin, Sec22b, Rab1, ERGIC-53 and Arf1 to the LCV (Derre and Isberg, 2004; Kagan and Roy, 2002; Kagan et al., 2004), which are needed for its replication. The mature LCV retains ER-like properties, including the lumenal proteins calnexin, BiP and calreticulin and does not acquire Golgi markers (Derre and Isberg, 2004; Kagan and Roy, 2002; Kagan et al., 2004; Treacy-Abarca and Mukherjee, 2015). Given that the LCV is derived from the ER, and that CHC22 localizes to a compartment emerging from the ER, we tested whether CHC22 associates with membranes involved in LCV formation. A549 human lung adenocarcinoma cells were transiently transfected to express GFP-tagged CHC22 or GFP-tagged CHC17, then infected with *L.p.*. CHC22, but not CHC17, was associated with the membranes surrounding the LCV (Fig. 5 A and B). Similar LCV co-localization with CHC22 was observed in untransfected cells infected with *L.p.* and immuno-stained for endogenous CHC22 or CHC17 (Fig. 5 C). CHC22 did not localize with the isogenic avirulent *L.p.* mutant *ΔdotA,* which still enters cells but lacks a functional secretion system and cannot secrete effectors to create an ER-like vacuole (Fig. 5 A and B). To further address whether CHC22 is involved in transfer of membrane to the LCV, we treated A549 cells with siRNA targeting CHC22 prior to infection. The resulting CHC22 down-regulation significantly compromised recruitment of Sec22b to the bacterial vacuole at 1 hour post-infection (Fig. S4 A and B), suggesting defective vacuole maturation. This was confirmed by assessing bacterial replication eight hours after infection with WT or *ΔdotA L.p.* strains following CHC22 or CHC17 depletion. CHC22 depletion reduced the proportion of vacuoles containing >4 *L.p.* by more than 9-fold, while CHC17 depletion reduced vacuoles with >4 *L.p.* by only 2-fold (Fig. 5 D), indicating that CHC22 is required to form a replicative vacuole and a possible role for CHC17 during bacterial uptake. The latter conclusion is supported by the observation that after CHC22 down-regulation, vacuoles with 1 *L.p.*, indicating bacterial entry, were observed, but vacuoles with 1 *L.p.* were less frequent in cells depleted for CHC17 infected with an equivalent number of bacteria. Infection of cells transfected with siRNA targeting CHC22 with an equivalent number of avirulent *ΔdotA L.p.* showed that these bacteria could also enter cells, with 94% vacuoles observed harboring only 1 *L.p.* (Fig. 5 D). These observations indicate that *L.p.* specifically co-opts CHC22 to acquire membrane derived from the early secretory pathway, which is needed for maturation of a replication-competent LCV, and suggest that *L.p.* effectors might interact with CHC22 or its partners.

**Figure 5:**
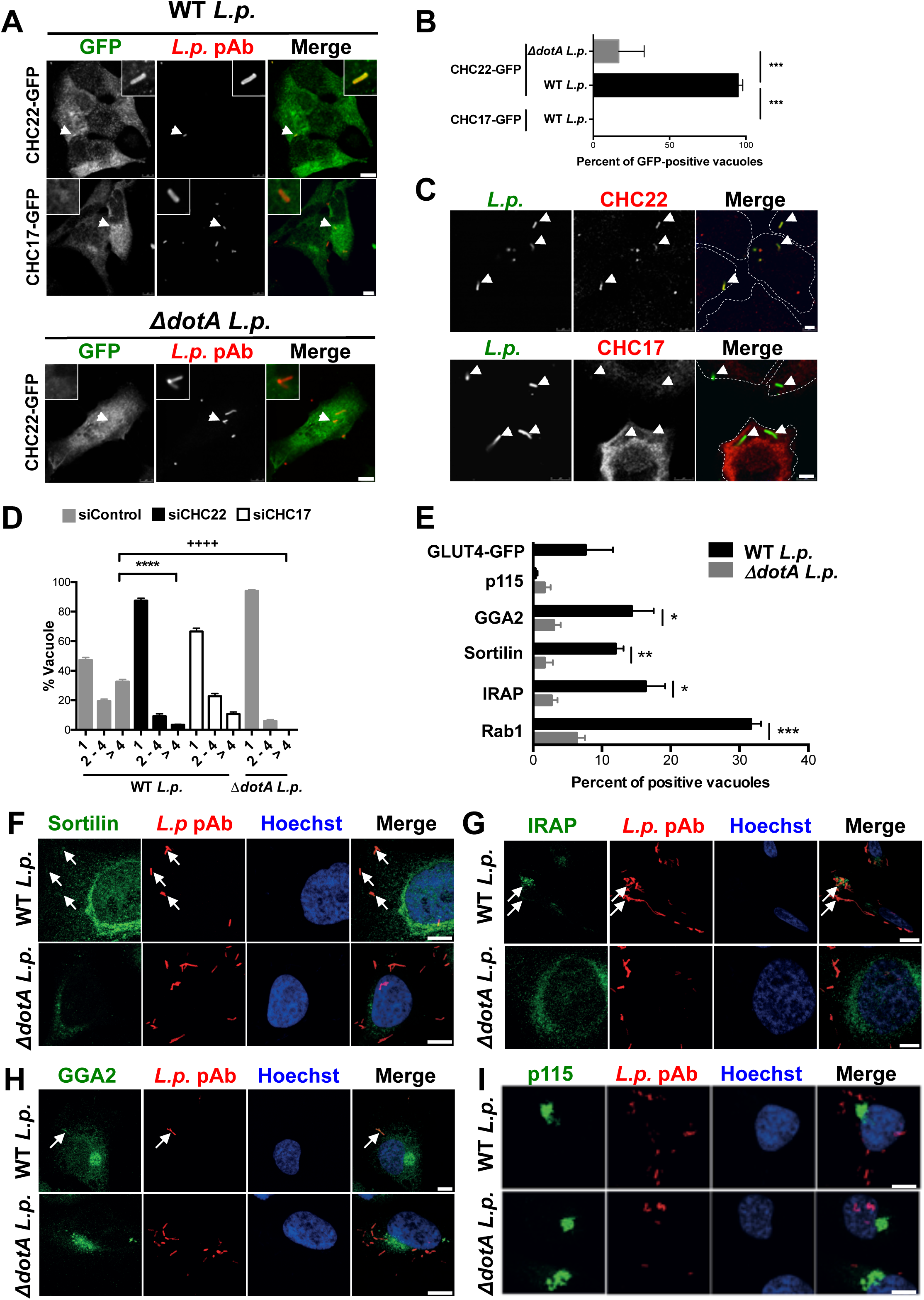
CHC22 and proteins involved in the GLUT4 pathway participate in membrane trafficking from the ERGIC. (A) Representative images of *Legionella pneumophila* (*L.p.*)-infected A549 cells transiently transfected with GFP-tagged CHC22 or CHC17 (green). One hour after infection with either wild type (WT) or mutant *ΔdotA L.p.* (MOI=50), bacteria were detected by immunofluorescence (IF, red). Arrows point to *L.p.* and boxed inserts (upper right or left) show *L.p.* region at 5X magnification. Scale bars: 10 µm for cells expressing CHC22-GFP and 7.5 µm for cells expressing CHC17-GFP. (B) Quantification of the proportion of *L.p.* vacuoles positive for CHC22 or CHC17. Data expressed as mean ± SEM, N=3, 4 to 35 vacuoles counted per experiment performed as represented in (A). One-way analysis of variance (ANOVA) followed by Bonferroni’s multiple comparison post-hoc test ***p<0.001. (C) Representative images of A549 cells infected with WT *L.p.* (MOI=50) immunolabeled for endogenous CHC22 or CHC17 (red) and *L.p.* (green) by IF. Arrows point to *L.p.*, dashed lines delineate cell borders. Scale bar: 5 µm. (D) Quantification of the proportion of replicative vacuoles (8 hours post-infection) containing 1, 2 to 4, or more than 4 WT or *ΔdotA L.p.* after treatment with siRNA targeting CHC22 (siCHC22) or CHC17 (siCHC17) or non-targeting siRNA (siControl). Data expressed as mean ± SEM, N=3, over 140 vacuoles counted per experiment. One-way ANOVA followed by Bonferroni’s multiple comparison post-hoc test was performed to compare the number of cells with a vacuole containing more than 4 bacteria. ****p<0.0001 versus siControl-transfected cells infected with WT *L.p.*. ++++p<0.0001 versus siControl-transfected cells infected with *ΔdotA L.p.* (E) Quantification of the proportion of *L.p.* vacuoles positive for GLUT4-GFP, p115, GGA2, sortilin, IRAP or Rab1 1h after infection with WT or *ΔdotA L.p.* in HeLa cells transiently expressing FcγRII (needed for *L.p.* infection) (Arasaki and Roy, 2010). Data expressed as mean ± SEM, N=3, 4 to 50 vacuoles counted per experiment. Two-tailed unpaired Student’s t-test with equal variances: *p<0.05, **p<0.01, ***p<0.001. (F-I) Representative immunofluorescence of HeLa cells one hour after infection with either wild type (WT) or mutant *ΔdotA L.p.* (MOI=50) stained for *L.p.* (red) and sortilin (F), IRAP (G), GGA2 (I), or p115 (green) (I). Hoechst stains the nuclei (blue). Arrows point to *L.p.*-containing vacuoles, Scale bars: 10 µm. Merged images in (A), (C), (F-I) show red/green overlap in yellow.

Since CHC22-mediated membrane traffic contributes to formation of the LCV, we addressed whether GLUT4 or other components of the GLUT4 trafficking pathway traffic to the LCV. After infection, we observed significant enrichment of sortilin, IRAP and GGA2 to the LCV, all known functional partners for intracellular GLUT4 sequestration (Li and Kandror, 2005; Shi et al., 2008; Shi and Kandror, 2005; Shi and Kandror, 2007; Watson et al., 2004) (Fig. 5 E-H). We also observed recruitment of Rab1, a host protein that is extensively modified by *L.p.* during infection (Mukherjee et al., 2011) (Figs. 5 E and S4 C). In contrast, the localization of GLUT4 (Figs. 5E and S4 D) or insulin effector Sec16a (Bruno et al., 2016) (Fig. S4 E and F) to the LCV was not statistically significant. We also did not detect p115 on the LCV (Fig. 5 E and I), confirming previous work from others suggesting that one of the *L.p.* effectors LidA bypasses the need for p115 by binding Rab 1 during host membrane recruitment (Derre and Isberg, 2004; Machner and Isberg, 2006). Thus, analysis of *L.p.* infection indicates that CHC22 actively traffics membranes from the early secretory pathway to the LCV, and that this pathway also traffics a subset of proteins involved in forming the GSC.

### CHC22 interacts with p115 and each influences stability of different partners for GLUT4 membrane traffic

Previous work implicated the vesicle tether p115 in murine GSC formation in 3T3-L1 adipocytes by showing an interaction between p115 and IRAP and demonstrating that expression of the interacting fragment of p115 had a dominant negative effect on the GLUT4 insulin response (Hosaka et al., 2005). Given the high degree of co-localization between CHC22 and p115 in human cells, we addressed the possibility that CHC22 and p115 might also associate. Endogenous CHC22 or CHC17 were immunoprecipitated from lysates of HeLa-GLUT4 (Fig. 6 A) or human skeletal muscle myotubes (Fig. 6 B) using an antibody highly specific for CHC22 and the most specific antibody available for CHC17, which has slight cross-reactivity with CHC22. An association between p115 and CHC22 was detected in both cell lysates and p115 was not co-immunoprecipitated with CHC17 (Fig. 6 A and B). As previously found (Vassilopoulos et al., 2009), GLUT4 co-immunoprecipitated preferentially with CHC22 compared to its association with CHC17 in both types of cells. We also previously showed that, compared to CHC17, CHC22 preferentially co-immunoprecipitated with the adaptor GGA2 (Dannhauser et al., 2017; Vassilopoulos et al., 2009), a reported interactor of sortilin, which is involved in retrograde transport of recaptured GLUT4 in murine adipocytes (Pan et al., 2019; Shi et al., 2008; Shi and Kandror, 2005; Shi and Kandror, 2007). Here we found that, in both HeLa-GLUT4 cells and human skeletal muscle myotubes, sortilin specifically co-immunoprecipitated with CHC22 but not CHC17 (Fig. 6 C and D). CHC22 was not isolated by the anti-sortilin monoclonal antibody used, suggesting that the relevant epitope of sortilin was not accessible when associated with CHC22. Thus CHC22 complexes with molecules from both the early secretory pathway and the GLUT4 retrograde transport pathway.

**Figure 6:**
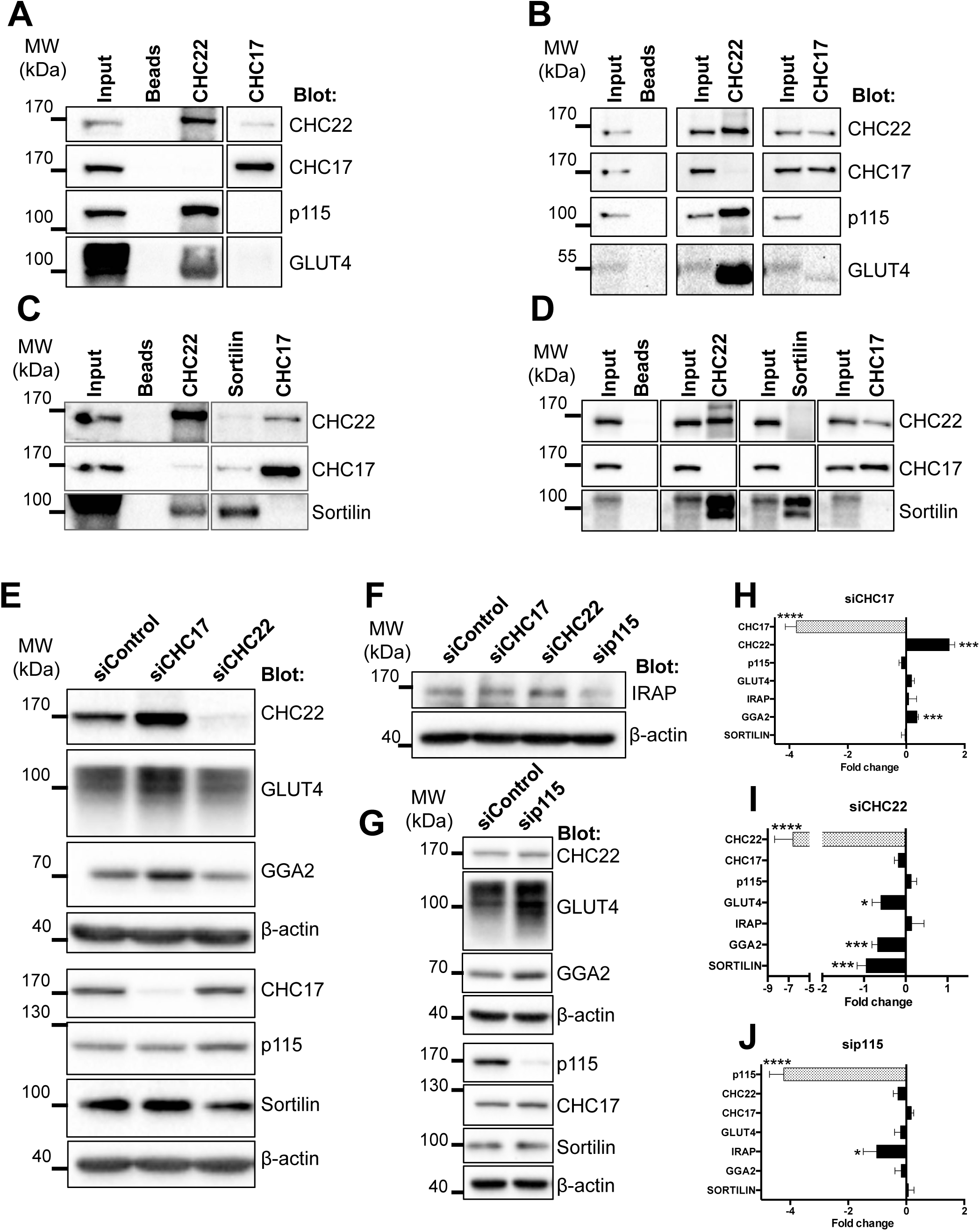
CHC22 interacts with p115 and each influences stability of different partners involved in GLUT4 membrane traffic. (A-D) Representative immunoblots of immunoprecipitates of CHC22, CHC17 (A-D) or Sortilin (C and D) from HeLa-GLUT4 cells (A and C) and hSKMC-AB1190 (B and D) immunoblotted for CHC22, CHC17, p115, GLUT4 and Sortilin. The position of molecular weight (MW) markers is indicated in kilodaltons (kDa). (E-G) Representative immunoblots of HeLa-GLUT4 cells transfected with siRNA targeting CHC22, CHC17 (E, F) or p115 (F, G) or with non-targeting control siRNA (40 nM for 72h) showing levels of CHC22, GLUT4, GGA2, CHC17, p115, sortilin and β-actin (E, G) or levels of IRAP and β-actin (F). The position of molecular weight (MW) markers is indicated in kilodaltons (kDa). (H-J) Quantifications of immunoblot signals as shown in (E-G). Blot signals were normalized to β-actin for each experiment and the fold change (negative values indicate decrease and positive values indicate increase) relative to the normalized signal in control siRNA-treated cell lysates is plotted. Data expressed as mean ± SEM, N=7-8. Two-tailed unpaired Student’s t-test, with Welch’s correction where variances were unequal: *p<0.05; ***p<0.001, ****p<0.0001.

To address the relationship of the CHC22-p115 complex with other components of the GLUT4 trafficking pathway, we assessed the effects of CHC22 and p115 depletion on each other and on GLUT4 traffic participants, compared to CHC17 depletion in HeLa-GLUT4 cells. In particular, we focused on the fate of IRAP, sortilin and GGA2. These proteins were all found associated with the LCV (Fig. 5 E-H) and all three have been implicated in rodent GSC formation, as well as found in complexes with each other (Li and Kandror, 2005; Shi et al., 2008; Shi and Kandror, 2005; Shi and Kandror, 2007; Watson et al., 2004). CHC22 depletion destabilized GLUT4, sortilin and GGA2 while p115 and IRAP remained unchanged (Fig. 6 E, F and I). Conversely, p115 depletion destabilized IRAP (Fig. 6 F and J) but none of the other components. None of the GLUT4 trafficking components were destabilized upon CHC17 depletion (Fig. 6 E, F and H), which, as previously observed (Dannhauser et al., 2017; Esk et al., 2010; Vassilopoulos et al., 2009), stabilized CHC22, likely due to increased membrane association as a result of reduced competition for shared adaptors such as GGA2 and AP1. Combining these observations with the immunoprecipitation results suggests one minimal complex between CHC22, GGA2, sortilin and GLUT4 and another minimal complex between CHC22, p115, IRAP and GLUT4, with the former playing a role in retrograde GLUT4 sorting and the latter playing a role in sorting newly synthesized GLUT4.

### Membrane traffic to the human GSC requires CHC22 and p115, but not GM130

To investigate the involvement of the early secretory pathway in sorting GLUT4 during GSC formation in human cells, we depleted CHC22, p115 or the cis-Golgi tether protein GM130 from HeLa-GLUT4 cells using siRNA (Fig. 7 A) and assessed the distribution of GLUT4 by confocal microscopy (Fig. 7 B and C). Depletion of p115 or of CHC22 caused loss of perinuclear GLUT4 and dispersion in the cell periphery, with down-regulation of one affecting the distribution of the other (Fig. 7 B). We did not detect any impact of GM130 depletion on GLUT4 subcellular distribution, though GM130 depletion did partially alter p115 distribution (Fig. 7 C). Since IRAP and sortilin were found to associate with p115 and CHC22, we also tested the effects of their down-regulation on GLUT4 distribution and observed no obvious effects of these treatments (Fig. 7 A, D, E), indicating differences in requirements for IRAP and sortilin in targeting GLUT4 in this human model compared to the murine adipocyte model.

**Figure 7:**
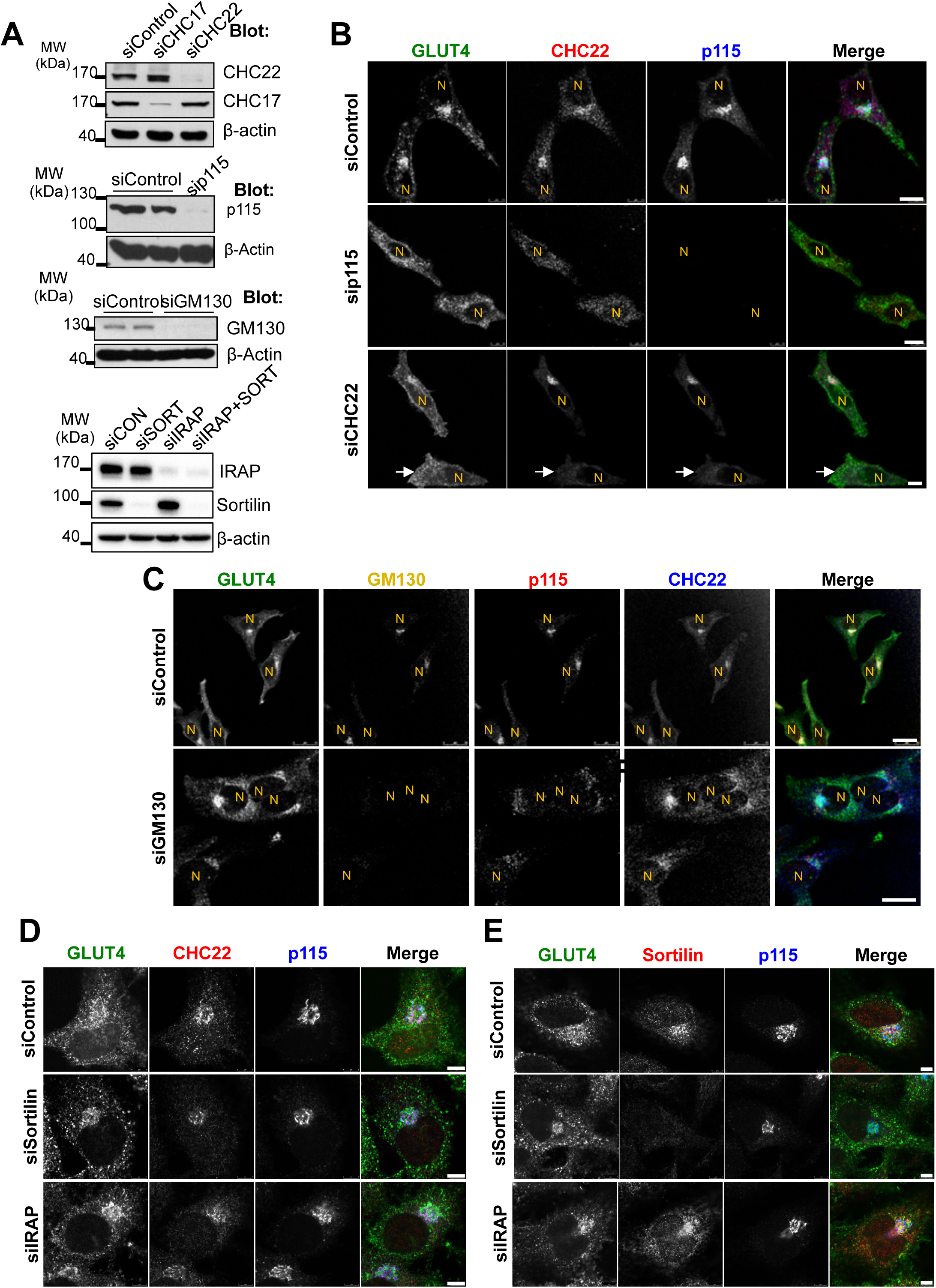
Depletion of CHC22 or p115, but not GM130, sortilin or IRAP disrupt perinuclear targeting of GLUT4. (A) Immunoblotting for CHC22, CHC17, p115, GM130 and β-actin after transfection of HeLa-GLUT4 cells with siRNA targeting CHC17, CHC22, p115, GM130, sortilin, IRAP or with non-targeting control siRNA. The position of molecular weight (MW) markers is indicated in kilodaltons (kDa). (B) Representative immunofluorescence (IF) staining for CHC22 (red) and p115 (blue) in HeLa-GLUT4 cells after siRNA transfection as in (A), with GLUT4 detected by GFP tag (green). N, nuclei. Arrow points to CHC22-depleted cell. Scale bars: 10 µm. (C) Representative IF staining for GM130 (yellow), p115 (red) and CHC22 (blue) in HeLa-GLUT4 cells after treatment with siRNA targeting GM130 or with control siRNA, with GLUT4 detected by GFP tag (green). Individual antibody staining is shown in black and white, while the merged image shows all four colors with overlaps in white. Scale bars: 25 µm. (D and E) Representative IF staining for GLUT4 (green), CHC22 or sortilin (red) and p115 (blue) in HeLa-GLUT4 cells after treatment with siRNA targeting sortilin or IRAP or with non-targeting control. Scale bars: 5 µm. Merged images show red/green overlap in yellow, red/blue overlap in magenta, green/blue overlap in turquoise, and red/green/blue overlap in white.

To determine the functional effects of altering GLUT4 distribution in the HeLa-GLUT4 cells, we evaluated how depletion of CHC22, p115, GM130, IRAP, and sortilin affected insulin-induced GLUT4 translocation, as assessed by FACS analysis (Fig. 8 A and B). Insulin-stimulated GLUT4 translocation was lost from cells depleted of p115 or CHC22 while CHC17 depletion had a partial effect on GLUT4 translocation (Fig. 8 A), consistent with previous observations for CHC22 and CHC17 down-regulation (Vassilopoulos et al., 2009). Corresponding to immunofluorescence analyses (Fig. 7 C-E), down-regulation of GM130, IRAP and sortilin (or the combination of IRAP and sortilin) did not affect insulin-stimulated GLUT4 translocation (Fig. 8 A and B). To confirm that GM130 depletion affected export from the Golgi, we demonstrated reduction of alkaline phosphatase secretion from the siRNA-treated cells (Tokumitsu and Fishman, 1983) (Fig. S5). These translocation assays further indicate that CHC22 and p115 are essential for formation of the human GSC and that this process requires membrane traffic from the early secretory pathway that bypasses the Golgi.

**Figure 8:**
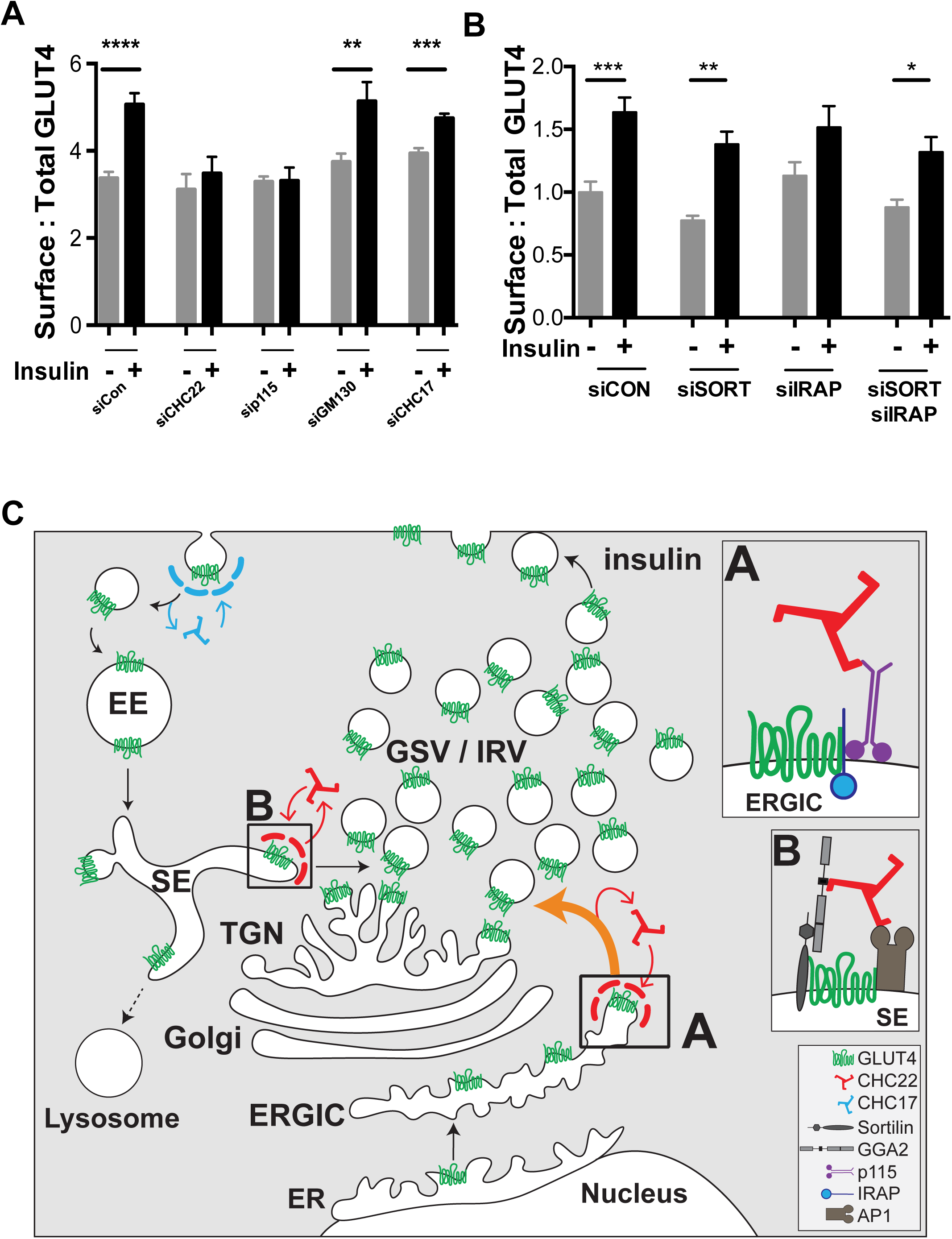
Formation of the human insulin-responsive GLUT4 pathway involves membrane traffic from the ERGIC and supports a model for two routes to GLUT4 sequestration. (A and B) Insulin-stimulated GLUT4 translocation in HeLa-GLUT4 cells as quantified by FACS analysis of surface:total GLUT4. Cells were pre-treated with siRNA targeting CHC22, CHC17, p115, GM130, sortilin, IRAP, sortilin plus IRAP, or with non-targeting control siRNA (siCon) as in Fig. 7 A, then incubated with (+) or without (-) insulin. For the experiments in (A), data is expressed as mean ± SEM, N=9, 10,000 cells acquired per experiment. One-way ANOVA followed by Bonferroni’s multiple comparison post-hoc test **p<0.01, ***p<0.001, ****p<0.0001 versus untreated (-). For the experiments in (B), data is expressed as mean ± SEM, N=7, 10,000 cells acquired per experiment. One-way ANOVA followed by Bonferroni’s multiple comparison post-hoc test *p<0.05, **p<0.01, ***p<0.001 versus untreated. (C) Proposed model for the roles of CHC22 in the human GLUT4 pathway. Newly synthesized GLUT4 traffics from the endoplasmic reticulum (ER) to the ER-to-Golgi Intermediate Compartment (ERGIC). At the ERGIC, a complex forms between IRAP and p115 that promotes binding of CHC22 clathrin and sequesters GLUT4 through its IRAP interaction (Shi et al., 2008) (box A). Formation of the CHC22 clathrin coat at the ERGIC then facilitates sorting of GLUT4 to the intracellular region where GLUT4 storage vesicles (GSV) and insulin responsive GLUT4 vesicles (IRV) are formed. After insulin-mediated GLUT4 translocation and GLUT4 re-uptake (by CHC17 clathrin), a complex forms (box B) between endosomal GLUT4, sortilin and the clathrin adaptor GGA2, which promotes CHC22 recruitment. Endosomal GLUT4 sorting also involves clathrin adaptor AP1 (Blot and McGraw, 2008; Gillingham et al., 1999), further participating in CHC22 recruitment to endosomes. Formation of the CHC22 coat on sorting endosomes facilitates GLUT4 traffic to the TGN via the retrograde pathway, enabling replenishment of the intracellular GSV/IRV pool.

## Discussion

CHC22 clathrin is required for formation of the insulin-responsive GLUT4 storage compartment (GSC) in human muscle and fat (Esk et al., 2010; Vassilopoulos et al., 2009) and has been implicated in specialized membrane traffic to dense core granules in neuronal cells (Nahorski et al., 2018). Both of these roles require diversion of intracellular cargo into privileged storage compartments so that the cargo is sequestered from degradation. Previous studies indicated a role for CHC22 clathrin in retrograde transport from endosomes (Esk et al., 2010), where CHC17 has been implicated in returning released GLUT4 to the GSC in murine cells (Gillingham et al., 1999; Li and Kandror, 2005). However, depletion of CHC22 from human muscle cells abrogates GSC formation even in the presence of CHC17 (Vassilopoulos et al., 2009), so we suspected a second pathway for CHC22 function in GLUT4 transport unique to this isoform of clathrin. We focused on pathways that would be involved in biosynthetic formation of the GSC, as evidence suggests that GLUT4 arrives at the GSC prior to its expression on the plasma membrane (Watson et al., 2004). Discovering strong co-localization of CHC22 with the ERGIC markers p115 and ERGIC-53, we investigated a role for CHC22 in formation of the replication vacuole of *Legionella pneumophila* (*L.p.*), which acquires membrane from the early secretory pathway to evade degradative compartments (Derre and Isberg, 2004; Kagan and Roy, 2002). We found that CHC22 was required for bacterial replication and formation of the *L.p.*-containing vacuole (LCV), and that components of the GLUT4 trafficking pathway localized to the LCV, though the presence of GLUT4 itself was highly variable. A difference between LCV and GSC formation is a requirement for cellular p115. We showed that p115 was needed for human GSC formation, and it has previously been implicated in murine GSC formation (Hosaka et al., 2005), but *L.p.* bacteria have an effector protein that replaces p115 (Machner and Isberg, 2006). Our studies therefore indicate that human GSC formation requires CHC22-dependent membrane derived from the ERGIC, in a pathway similar to that co-opted by *L.p.* bacteria.

Tracking newly synthesized GLUT4, using the RUSH system, we observed that GLUT4 resides for a longer period of time with markers of the early secretory pathway compared to the behavior of constitutively secreted GLUT1 after their release from the ER. We also found that while formation of the human GSC was sensitive to depletion of p115 and CHC22, GSC formation was not affected by depletion of GM130. These data suggest that the CHC22-mediated pathway transports GLUT4 to a site where it can be sequestered for generation of insulin-responsive vesicles and further suggests a Golgi bypass occurs in this step of GSC biogenesis. The reported slow maturation of carbohydrate side chains on GLUT4 compared to GLUT1 is consistent with the proposed Golgi bypass (Hresko et al., 1994; Hudson et al., 1992). Delayed modification of GLUT4 carbohydrate could result from pre-Golgi diversion of most newly synthesized GLUT4 from the ERGIC, followed by a process of carbohydrate maturation that depends on GLUT4 secretion and recapture through a retrograde pathway (Shewan et al., 2003) where low-level carbohydrate modification can occur by Golgi enzymes being recycled to their home compartments (Fisher and Ungar, 2016). In our studies, we localized ERGIC membrane with CHC22 and/or p115 in close proximity to, but not completely overlapping with, compartments containing GLUT4 internalized after insulin-mediated release and intracellular compartments marked by STX-6, a TGN marker that co-localizes with internalized GLUT4. This is consistent with a biogenesis pathway connecting GLUT4 emerging from the ERGIC with the pool of GLUT4 sequestered after internalization for reformation of insulin-responsive vesicles. It is also consistent with ER/ERGIC-associated proteins being involved in insulin-mediated GLUT4 release, such as the TUG vesicle tethering protein (Orme and Bogan, 2012) and the ER-exit site protein Sec16A (Bruno et al., 2016). Proximity of GLUT4 emerging from early secretory compartments with compartments generating insulin-responsive GLUT4 vesicles could explain sharing of effector molecules and would also explain the general insensitivity of the GSC to BFA disruption, as shown here and earlier (Martin et al., 2000).

Our demonstration that CHC22 functions in transport from the early secretory pathway defines a membrane traffic step in which the canonical CHC17 clathrin is not involved (Brodsky, 2012). We show here that CHC22 co-immunoprecipitates with p115 and sortilin and that CHC17 does not, further demonstrating that the two clathrins form distinct complexes (Vassilopoulos et al., 2009), localize to distinct cellular regions (Liu et al., 2001) and form distinct coated vesicles (Dannhauser et al., 2017). It was previously shown that p115 interacts with IRAP (Hosaka et al., 2005), a protein that binds GLUT4 and is co-sequestered in GLUT4 vesicles (Shi et al., 2008) and that expression of a p115 fragment prevents GSC formation in murine cells (Hosaka et al., 2005). We show here that p115 down-regulation reduces the stability of IRAP. Thus, we propose that CHC22-p115-IRAP interaction occurs in the human ERGIC and triggers the coalescence of a protein domain that captures GLUT4 for sorting to the GSC (Fig. 8C). Taking into account our earlier demonstration of a role for CHC22 in retrograde transport from endosomes and its preferential interaction with the endosome-TGN adaptor GGA2 (Dannhauser et al., 2017; Vassilopoulos et al., 2009) as well as the interaction with sortilin shown here, we propose that a second complex involving CHC22, sortilin and GGA2 sorts internalized GLUT4 to the compartment where insulin-responsive vesicles are generated (Fig. 8C). Thus, CHC22 clathrin can play a role in sorting both newly synthesized and internalized GLUT4 to the human GSC.

GLUT4 membrane traffic has primarily been studied using murine adipocyte and rat myoblast cell lines, which do not express CHC22 as a result of gene loss in the rodent lineage (Fumagalli et al., 2019; Wakeham et al., 2005). Such studies have established that the major pathway for targeting GLUT4 to the GSC in rodent cells relies on retrograde transport of GLUT4, via endosomal-TGN sorting, after its release to the cell surface from the GSC and uptake by CHC17 (Bryant et al., 2002; Jaldin-Fincati et al., 2017). Our studies here (Fig. 1) support the existence of this retrograde pathway in human cells. It is also reported that in rodent cells GLUT4 reaches the GSC prior to its insulin-stimulated release to the cell surface (Lamb et al., 2010; Watson et al., 2004), and in rodent cells there is involvement of p115 in GSC formation (Hosaka et al., 2005). Thus, both direct targeting and endocytic recapture pathways seem to be involved in GSC formation in humans and rodents. In humans the CHC22 coat mediates both pathways, but rodents only have the CHC17 coat to mediate retrograde sorting. So rodents must rely simply on coalescence of relevant cargo (GLUT4, IRAP and p115) in the ERGIC to segregate them from other proteins constitutively leaving the secretory pathway and this is likely less efficient, perhaps with some GLUT4 directly accessing the cell surface for rapid re-uptake in the absence of insulin (Martin et al., 2000). In this case, population of the rodent GSC with GLUT4 is mainly a result of the retrograde recycling pathway, in which sortilin and IRAP have been shown to participate (Jordens et al., 2010; Pan et al., 2019; Pan et al., 2017; Sadler et al., 2019). In the case of human cells, CHC22 can actively capture GLUT4 and partners for diversion to the GSC, providing a more robust route to GSC formation following biosynthesis, with the GSC also replenished with GLUT4 by the endocytic-retrograde recycling pathway. The fact that humans have a very stable coat contributing to each sorting step (CHC22 is more stable than CHC17) (Dannhauser et al., 2017; Liu et al., 2001), may explain why neither sortilin nor IRAP knockdown affected GSC formation in our human model, even if they are functionally important cargo for GLUT4 sorting (Fig. 7E and F). The species difference in membrane dynamics of GLUT4 traffic has the effect that human cells cannot form a GSC with only CHC17 and require both CHC22 pathways. However, the presence of CHC22 enhancing biosynthetic GSC formation may have the consequence that humans are able to sequester intracellular GLUT4 more efficiently than species without CHC22, a trait that may contribute to a tendency to insulin resistance.

Identification of this sorting pathway for GLUT4 from the ERGIC to the GSC adds to the variety of sorting pathways that are known to emerge from the ERGIC, sustaining both conventional (ER-to-Golgi) (Kondylis and Rabouille, 2003; Sohda et al., 2007) and unconventional (Golgi bypass) pathways for autophagy (Ge et al., 2013; Ge and Schekman, 2014) or cargo exocytosis (Piao et al., 2017). In humans and even in CHC22 transgenic mice, CHC22 expression parallels that of GLUT4, with its highest expression in GLUT4-expressing tissues (Hoshino et al., 2013). However, unlike the tight regulation of GLUT4 expression, CHC22 is expressed at low levels in additional cell types (Nahorski et al., 2015). Thus, the CHC22 sorting pathway emerging from the ERGIC that we define here and CHC22-mediated retrograde sorting may also operate in tissues that do not express GLUT4 to target yet-unidentified cargo to specialized organelles, avoiding the conventional secretory or endocytic pathways.

## Materials and Methods

### Plasmids

The HA-GLUT4-GFP construct was a gift from Dr Tim McGraw (Lampson et al., 2000). The plasmid encoding human GLUT1 was from OriGene. The haemagglutinin (HA-)tag sequence atcgattatccttatgatgttcctgattatgctgag was inserted at base pair 201 (between amino acids 67 and 68 of the exofacial loop of GLUT1) using the Q5 site-directed mutagenesis kit from New England Biolabs (NEB, USA). HA-GLUT4 and HA-GLUT1 were extracted using AcsI and EcoRI restriction enzymes and Cutsmart buffer from NEB and the agarose gel extraction kit from Qiagen. The inserts were then ligated into the RUSH plasmid containing the ER Ii-hook fused to streptavidin, to generate the HA-GLUT4-SBP-GFP construct (Boncompain et al., 2012; Boncompain and Perez, 2013). In order to generate the HA-GLUT1-SBP-mCherry plasmid, we swapped the GFP tag in the HA-GLUT1-SBP-GFP for mCherry, using the SbfI and FseI restriction enzymes. The generation of plasmids encoding GFP-tagged CHC22 and CHC17 has been described elsewhere (Esk et al., 2010).

### Cell culture

All cell lines were maintained at 37°C in a 5% CO_2_ atmosphere. The HeLa cell line stably expressing GLUT4 (HeLa-GLUT4) was generated by transfection of HeLa cells with the plasmid encoding HA-GLUT4-GFP (Dawson et al., 2001; Lampson et al., 2000; Quon et al., 1994). Transfectants were selected in growth medium supplemented with 700 µg/mL G418 then maintained in growth medium with 500 µg/mL G418. The human skeletal muscle cell line LHCNM2 was described elsewhere (Esk et al., 2010; Vassilopoulos et al., 2009; Zhu et al., 2007). A549 human lung carcinoma cells were obtained from the ATCC. HeLa and A549 cells were grown in Dulbecco’s Modified Eagle Medium high glucose supplemented with 10% FBS (Gibco), 50 U/mL penicillin, 50 µg/mL streptomycin (Gibco), 10 mM Hepes (Gibco). LHCNM2 cells were grown in proliferation medium: DMEM MegaCell (Sigma) supplemented with 5% FBS (Gibco), 2 mM L-Glutamine (Sigma), 1% non-essential amino acids (Sigma), 0.05 mM β-mercaptoethanol (Gibco) and 5 ng/mL bFGF (Thermo Fisher). When full confluency was reached, cells were switched to differentiation medium: DMEM (Sigma) supplemented with 2 mM L-Glutamine (Sigma), 100 IU penicillin and 100 µg/mL streptomycin (Gibco). The human myoblast cell line AB1190 was immortalized at the platform for immortalization of human cells from the Institut de Myologie (Paris). These cells were grown in complete Skeletal Muscle Cell Growth Medium (Promocell) supplemented with serum to reach 20% final concentration (V/V). These cells were transfected to express HA-GLUT4-GFP and permanently transfected myoblasts (hSkMC-AB1190-GLUT4) were selected for their ability to differentiate into myotubes. Differentiation of confluent hSkMC-AB1190 myoblasts and hSkMC-AB1190-GLUT4 myoblasts was induced by incubating the cells in differentiation medium for 6 to 7 days: DMEM (Gibco), Gentamycin 50 µg/ml (Gibco) + insulin 10 µg/ml (Sigma). All cell lines used were tested negative for mycoplasma infection.

### Small RNA interference

Targeting siRNA was produced (Qiagen) to interact with DNA sequences AAGCAATGAGCTGTTTGAAGA for CHC17 (Esk et al., 2010), TCGGGCAAATGTGCCAAGCAA and AACTGGGAGGATCTAGTTAAA for CHC22 (1:1 mixture of siRNAs were used)(Vassilopoulos et al., 2009) and AAGACCGGCAATTGTAGTACT for p115 (Puthenveedu and Linstedt, 2004). Targeting siRNA against sortilin and IRAP were purchased from OriGene (SR304211, SR302711, respectively). Non-targeting control siRNA was the Allstars Negative Control siRNA (Qiagen). siRNA targeting GM130 and scrambled negative control siRNA were purchased from OriGene. For siRNA treatments, cells were seeded (10,000 cells/cm^2^) in 6- or 24-well plates in culture medium. The next day, the cells were transfected with siRNAs complexed with JetPrime (PolyPlus). For targeting CHC17, p115 and GM130, 20 nM of siRNA was used per treatment. For targeting CHC22, 20 nM of siRNA was used per treatment, except for Western blot experiments in Figure 6 where 40 nM of siRNA were used. For IRAP and sortilin knockdowns, 30 and 40 nM of siRNA were transfected, respectively. Six hours after siRNA transfection, cells were returned to normal growth conditions and then harvested for analysis or fixed for imaging 72 h later. Silencing was assessed by immunoblotting.

### Transfection

For transient DNA transfection, cells were seeded (21,000 cells/cm^2^) in 24-well plates in culture medium. The next day, the cells were transfected with plasmid DNA complexed with JetPrime (PolyPlus) in a 1:2 mixture (DNA/JetPrime). For RUSH experiments, 0.5 μg of DNA was used. 0.25 μg of DNA was used for all other experiments. 6 h after DNA transfection, cells were returned to normal growth conditions and then fixed for imaging 24 h later.

### Antibodies and reagents

Primary antibody concentrations ranged from 1-5 µg/mL for immunoblotting and IF assays. Mouse monoclonal anti-CHC17 antibodies (TD.1 (Nathke et al., 1992) and X22 (Brodsky, 1985)), and affinity-purified rabbit polyclonal antibody specific for CHC22 and not CHC17 (Vassilopoulos et al., 2009) were produced in the Brodsky laboratory. Mouse monoclonal anti-p115 antibody (clone 7D1) has been described (Waters et al., 1992). Mouse monoclonal anti-GGA2 was a gift from Dr Juan Bonifacino (US National Institutes of Health, Bethesda, MD, USA). Rabbit polyclonal antibody anti-*L.p.* was a gift from Dr Craig Roy (Yale University, New Haven, CT, USA). Mouse monoclonal antibody anti-*L.p.* was made in the Mukherjee lab. Commercial sources of antibodies were as follows: Rabbit polyclonal anti-CHC17 (Abcam), rabbit polyclonal anti-CHC22 antibody (Proteintech), mouse monoclonal anti-β-COP (clone maD, Sigma), rabbit polyclonal anti-IRAP (#3808, Cell Signaling Technology), rabbit monoclonal anti-IRAP (clone D7C5, #6918, Cell Signaling Technology), rabbit polyclonal anti-phospho AKT Ser473 (#9271, Cell Signaling Technology), rabbit polyclonal anti-phospho-AS160 Thr642 (#4288, Cell Signaling Technology), rabbit polyclonal anti-AS160 (#2447, Cell Signaling Technology), goat polyclonal anti-GLUT4 (C-20, Santa-Cruz Biotechnologies), Rabbit anti-GLUT4 (Synaptic Systems), mouse monoclonal anti-calreticulin (clone FMC75, Stressgen Bioreagents), sheep polyclonal anti-TGN46 (AHP500G, Biorad), goat polyclonal anti-GM130 (P-20, Santa-Cruz Biotechnologies), sheep polyclonal anti-Sec22b (AHP500G, Creative Diagnostics), mouse monoclonal anti-ERGIC-53 (clone 2B10, OriGene), rabbit polyclonal anti-ERGIC-53 (E1031, Sigma), rabbit monoclonal anti-LMAN1 (clone EPR6979, Abcam), rabbit polyclonal anti-sortilin (ab16640, Abcam), Rabbit polyclonal anti-sortilin (12369-1-AP, Proteintech), mouse monoclonal anti-sortilin (clone EPR15010, ab188586, Abcam), mouse monoclonal anti-STX-6 (clone 30/Syntaxin 6, Becton Dickinson), mouse monoclonal anti-β actin (clone AC-15, Sigma), mouse monoclonal anti-HA (clone 16B12, Covance), mouse monoclonal anti-MHCI W6/32 (produced from the hybridoma in the Brodsky lab) has been described (Barnstable et al., 1978), goat polyclonal anti-Rab1 (orb153345, Biorbyt), chicken anti-GFP (A10262, Invitrogen). The commercial anti-CHC22 from Proteintech was confirmed in our laboratory to be specific for CHC22 and not CHC17 (Fig. S1 A). For IF, secondary antibodies coupled to FITC, Alexa Fluor 488, Alexa Fluor 555, Alexa Fluor 562 or to Alexa Fluor 647 (Thermo Fisher) were used at 1:500. For Western blotting, antibodies coupled to HRP (Thermo Fisher, Biorad) were used at 1:10,000. Brefeldin A (BFA) was from Sigma.

### Legionella pneumophila

WT and *ΔdotA Legionella* strains were gifts from Dr Craig Roy’s (Yale University). The parental strain (wild type) was *L.p.* serogroup 1 strain L.p.01, and the variant strain *ΔdotA* were isogenic mutants described previously (Berger et al., 1994; Zuckman et al., 1999). Single colonies of *L.p.* were isolated from charcoal yeast extract plates after growth for 2 days at 37°C. DsRed-expressing WT *L.p.* were grown on charcoal yeast extract plates containing 500 μM isopropyl+- o-thiogalactopyranoside (IPTG) for 2 days at 37°C to induce expression of the fluorescent protein.

### Infection and analysis of replicative vacuoles

A549 cells were seeded 10^5^ cells per 2 cm^2^ on coverslips. Cells were infected at a multiplicity of infection (MOI) of 25 with WT or *ΔdotA L.p.* strains. Immediately after adding *L.p.* to the medium, cells were centrifuged at 400 × g for 15 min, then left at 37°C for an additional 45 min. Cells were then washed 3X with PBS and incubated in growth medium for the indicated time. To analyse replicative vacuoles, cells were directly infected with WT or *ΔdotA L.p.* at a MOI of 50 for 1h for labelling with antibodies or transfected with siRNA (20 nM) or with plasmids encoding HA-GLUT4-GFP, CHC22-GFP or CHC17-GFP 72h before infection. Infected cells (transfected or not transfected) were incubated for 8h post-infection, then washed 3X with PBS, fixed with 2.5% paraformaldehyde (PFA) for 30 min and labelled with antibody to detect bacteria for counting the number per replicative vacuole and with antibodies to identify compartment markers.

### Immunofluorescence

Cells grown on 1.5# glass coverslips (Warner Instruments) were washed (PBS, 4°C), fixed (2-4% PFA, 30 min, 4°C), permeabilized and blocked (PBS 0.5% saponin, 2% bovine serum albumin) for 1 hour at room temperature (RT). Cells were then incubated with primary antibodies (overnight, 4°C), washed (5X, PBS, 4°C) and incubated with species-specific secondary antibodies coupled to fluorophores (Thermo Fisher). Cells were then washed (5X, PBS, 4°C) and coverslips mounted on microscope slides using Prolong Antifade Diamond kit (Thermo Fisher). Samples were imaged using a Leica TCS SP8 inverted laser scanning confocal microscope equipped with two high sensitivity (HyD) detector channels and one PMT detector channel, a 63X (1.40 NA) HC Plan-Apo CS2 oil-immersion objective and five laser lines. Dyes were sequentially excited at 405 nm (DAPI), 488 nm (GFP, Alexa Fluor 488), 543 nm (Alexa 555), 561 nm (Alexa 568), and 633 nm (Alexa 647). Multicolor images (1024 × 1024 pixels) were saved as TIFF files in Leica LAS X Software (Leica) and input levels were adjusted using ImageJ (US National Institutes of Health, NIH). Labelling detected in individual channels is shown in black and white in figure panels. Merged images are presented in pseudo-color as described in the legends. Image quantification was performed using ImageJ. For each cell, individual marker fluorescence was measured in separate channels, and signals were adjusted to their dynamic ranges. Degree of marker overlap in individual cells was determined by Pearson’s correlation coefficients.

### Structured Illumination Microscopy (SIM)

Sample preparation (fixation and staining) steps were identical to confocal microscopy. Sample acquisition was performed on a Zeiss Elyra PS.1 microscope (Axio Observer.Z1 SR, inverted, motorized) through a 100X alpha Plan-Apochromat DIC M27 Elyra lens (oil-immersion, 1.46 NA). Fluorophore excitation was performed with a 50 mW HR diode emitting at 350 nm (BP 420-480/LP 750 filter), and a 200 mW HR diode emitting at 488 nm (BP 495-550/LP 750 filter), a 200 mW HR Diode Pumped Solid State laser emitting at 561 nm (BP 470-620/LP 750 filter) and a 260 mW HR diode emitting at 642 nm (LP 655 filter). Acquisition was performed using a pco.egde sCMOS camera and post-acquisition processing (channel alignment) was performed on the ZEN Black software Version 11.0.2.190.

### GLUT4 internalization experiments

The GLUT4 internalization protocol was adapted from previous work (Foley and Klip, 2014). HeLa-GLUT4 cells were seeded on coverslips in 24-well plate and grown to 80% confluency. On the day of the experiment, cells were washed (3X, PBS, 37°C) and serum starved 2 hours. Cell surface HA-GLUT4-GFP was labelled on ice for 30 min with mouse monoclonal anti-HA antibody. After washing (5X, PBS, 4°C), cells were placed in serum-free medium (37°C) for indicated times. Cells were then washed (3X, PBS, 4°C), fixed and processed for immunofluorescence detection of internalised anti-HA antibody.

### GLUT4 translocation assay using flow cytometry

HeLa-GLUT4 cells were seeded in 6-well plates and grown to 95% confluency. On the day of experiment, cells were washed (3X, PBS, 37°C), serum-starved (2 hours), then treated with insulin to a final concentration of 170 nM or the same volume of vehicle (water) diluted in serum-free medium for 15 minutes, 37°C. Cells were then placed on ice and rapidly washed (3X, PBS, 4°C) and fixed (PFA 2.5%, 30 min). After fixation, cells were washed (3X, PBS, RT) then blocked for 1 hour (PBS 2% BSA, RT) before incubation with monoclonal anti-HA antibody (45 min, RT) to detect surface GLUT4. After incubation, cells were washed (5X, PBS, RT) and incubated with Alexa Fluor 647-anti-mouse Ig (45 min, RT). Cells were then washed (5X, PBS, RT), gently lifted using a cell scraper (Corning), pelleted (200xg, 10 min) and re-suspended (PBS, 2% BSA, 4°C). Data was acquired with Diva acquisition software by LSRII flow cytometer (Becton Dickinson) equipped with violet (405 nm), blue (488 nm) and red (633 nm) lasers. Typically, 10,000 events were acquired and Mean Fluorescence Intensity (MFI) values for surface GLUT4 (Alexa Fluor 647) and total GLUT4 (GFP) were recorded using 660/20 and 530/30 filters, respectively. Post-acquisition analysis was performed using FlowJo software (Treestar) where debris were removed by FSC/SSC light scatter gating then fluorescence histograms were analyzed. The ratio of surface to total MFI was calculated to quantify the extent of GLUT4 translocation.

### Translocation assay using immunofluorescence

To test the effect of insulin and CHC22 depletion on the surface expression of endogenous Class I MHC molecules, cells were treated with siRNA targeting CHC22 or with control siRNA (as elsewhere in the Methods) then serum starved (1h, 37°C) before insulin stimulation (15 minutes, 170 nM, 37°C). Cells were then placed on ice, washed (PBS, 4°C) and incubated with primary antibodies (1h, 4°C), then washed (PBS, 4°C), fixed (2.5% PFA, 30 min, 4°C), washed (PBS, 4°C) and incubated with fluorescent secondary antibodies (30 min, 4°C). Cells were then washed (5X, PBS, 4°C) and coverslips were mounted on microscope slides using Prolong Antifade Diamond mounting medium (Thermo Fisher).

### RUSH assay

HeLa cells were seeded at 30,000 cells/cm^2^ on coverslips. The next day, cells were transfected with 0.5 μg of HA-GLUT1-SBP-GFP or HA-GLUT4-SBP-GFP plasmids for 6h, then switched to fresh medium. The next day, cells were treated with 40 μM biotin for the indicated times. Cells were then quickly placed on ice, washed (3X, PBS, 4°C), fixed (2.5% PFA, RT) and processed for immunofluorescence. For videomicroscopy acquisitions, cells were transfected simultaneously with HA-GLUT1-SBP-mCherry (2 µg) and HA-GLUT4-SBP-GFP (2 µg) following protocol described elsewhere (Jordan et al., 1996) and imaged on a spinning disk confocal microscope.

### Alkaline phosphatase secretion assay

HeLa-GLUT4 cells were seeded in 96-well plates, grown to 80% confluency and were transfected the next day with 20 nM targeting or control siRNA. After 48h, cells were transfected with the plasmid encoding secreted alkaline phosphatase. After 24h, fresh medium was added to the culture and 8h later, the media were harvested and the cells lysed. Alkaline phosphatase activity in the harvested medium and cell lysate was assessed using the Phospha-Light System kit (Applied Biosystems), following the manufacturer’s instructions and detected with a luminometer (Varioskan LUX multimode multiplate reader, Thermo Fisher scientific). The alkaline phosphatase secretion index was determined by calculating the ratio of alkaline phosphatase activity detected in the medium (secreted) to total alkaline phosphatase activity in the culture (medium plus cell lysate activity).

### Brefeldin A treatment

HeLa-GLUT4 or LHCNM2 myoblasts were grown on coverslips and exposed to BFA (10 µg/mL, 1h, 37°C) or vehicle (DMSO) in starvation medium (DMEM only). During the last 15 minutes, cells were incubated with insulin (170 nM, 15 minutes) or vehicle (water). Cells were then washed (3X, PBS, RT) and processed for immunofluorescence.

### Preparation of clathrin-coated vesicles (for anti-CHC22 characterization)

Clathrin coated vesicles (CCV) preparation was adapted from Keen et al. (Keen et al., 1979). Briefly, pig brains were blended in buffer A (100mM MES, 1mM EDTA, 0.5 mM MgCl2) supplemented with 0.5 mM PMSF. The preparation was centrifuged at 8,000 rpm (JA-17 rotor, Beckman) at 4°C for 30 min, then the supernatant was filtered to remove particles and centrifuged at 40,000 rpm (45 Ti rotor, Beckman) at 4°C for 60 min to pellet the CCVs. A small volume of buffer A supplemented with 0.02 mM PMSF was added to the CCV pellets before homogenization with a potter S homogenizer. A solution of 12.5% Ficoll 12.5% sucrose was added 1:1 to the CCV suspension and gently mixed. The CCV preparation was then centrifuged at 15,000 rpm (JA-17 rotor, Beckman) at 4°C for 40 min. The supernatant was collected, diluted 5-fold in buffer A supplemented with 1 mM phenylmethane sulfonyl fluoride (PMSF) and centrifuged 40,000 rpm (45 Ti rotor, Beckman) for 60 min at 4°C to pellet vesicles. The pellet was resuspended in buffer A for Tris extraction. Finally, the preparation was purified by gel filtration (Superose 6, GE Life Science). CCV were stored at −80°C in 10 mM Tris–HCl, pH 8.0.

### Purification of hub CHC22 (for anti-CHC22 characterization)

Hub CHC22 fragment was produced in BL21(DE3) bacteria (Novagen) by induction with 1 mM IPTG for 24 hr at 12°C. Bacterial pellets were resuspended in LysI (1M NaCl pH=8, 20 mM imidazole in PBS) supplemented with 1 mM PMSF, protease inhibitors (1 tab/10 mL, Roche), 40 µg/mL lysozyme, 0.1% β-mercaptoethanol (Sigma). Then LysII (1M NaCl, 0.5M guanidine HCl, 0.4% Triton X100 in PBS) was added at 1:1.25 (LysI:LysII) ratio and samples were spun at 40,000 rpm for 30 min at 4°C. Supernatant was ran through a Ni+ affinity NTA column, then washed with LysI and eluted with 1M NaCl pH8, 0.5M imidazole in PBS.

### Immunoblotting

Protein extracts from cells were quantified by BCA (Pierce), separated by SDS-PAGE (10% acrylamide), transferred to nitrocellulose membrane (0.2 µm, Biorad) and labelled with primary antibodies (1-5 µg/mL), washed and labelled with species-specific horseradish peroxidase-conjugated secondary antibodies (Thermo Fisher). Peroxidase activity was detected using Western Lightning Chemiluminescence Reagent (GE Healthcare). The molecular migration position of transferred proteins was compared to the PageRuler Prestain Protein Ladder 10 to 170 kDa (Thermo Fisher Scientific). Signal quantification was performed using Image J software (NIH).

### Immunoprecipitation

Confluent cells from a 500 cm^2^ plate were scraped off the plate, washed in ice-cold PBS and pelleted (300 g, 8 min, 4°C). The pellets were resuspended in ice-cold lysis buffer (NaCl 150 mM, HEPES 20 mM, EDTA 1 mM, EGTA 1 mM, Glycerol 10% (V/V), NP-40 0.25% (V/V)) supplemented with protease (1 tab/10mL, Roche) and phosphatase (Na_4_VO_3_ 2 mM) inhibitors. Cell suspensions were mechanically sheared (over 25 passages through a 27G needle), sonicated and centrifuged (500 g for 10 min, 4°C) to remove nuclei. Five to 10 µg of specific anti-CHC22 (Proteintech) and CHC17 (X22) antibodies were incubated with 7 mg of pre-cleared post-nuclear supernatants (overnight, 4°C). The samples were then incubated with washed protein G sepharose (PGS, 25 µL, GE Healthcare) for 1h (4°C) before three consecutive washing steps in lysis buffer. Pelleted PGS were resuspended in 30 µL of 1X Laemmli sample buffer and subjected to SDS-PAGE and immunoblotting. Species-specific HRP-conjugated Trueblot secondary antibodies (Rockland) were used for immunoblotting IP experiments.

### Statistical analyses

All calculations and graphs were performed with Microsoft Excel and GraphPad Prism softwares. P-values were calculated using unpaired two-tailed Student’s t-tests or two-way ANOVA followed by Tukey, Dunnett, Bonferroni or Sidak’s multiple comparisons test. Detailed statistical information including statistical test used, number of independent experiments, *p* values, definition of error bars is listed in individual figure legends. All experiments were performed at least three times, except for the immunoblots shown in Figs. 1 B and 7 A and the infection experiment shown in Fig. S4 D, which were performed twice. Immunofluorescence stainings showed in Figs. S3 A, B were performed once.

## Summary of Supplemental material

This manuscript contains 2 videos and 5 supplemental figures (Fig. S1-S5).

## Acknowledgments

This work was supported (sequentially) by grants to FMB from the National Institutes of Health USA (DK095663), Wellcome Trust (107858/Z/15/Z) and Medical Research Council (MR/S008144/1). Early work on this project was supported by a postdoctoral fellowship from the American Heart Association (13POST17180010) to SMC. SM is supported by NIH RO1 AI118974 and a grant from the Pew Charitable Trust (A129837). GWG and NJB acknowledge the support of Diabetes UK (grant 11/0004289). DK was partially funded for this work by The Wellcome Trust [ref: 204829] through the Centre for Future Health (CFH) at the University of York. FP and GB are funded by the Centre National de la Recherche Scientifique, the Fondation pour la Recherche Médicale (FRM DEQ20120323723), the Agence Nationale de la Recherche (ANR-12-BSV2-0003-01) and are part of Labex CelTisPhyBio (11-LBX-0038) and Idex Paris Sciences et Lettres (ANR-10-IDEX-0001-02 PSL). We are grateful to UCL Excellence Fellow Philip Dannhauser for clathrin protein samples for immunoblotting and for providing the protocols for their production. We thank the platform for immortalization of human cells from the Institut de Myologie for providing the immortalized human myoblast cell line hSkMC-AB1190. Cell culture studies reported here utilize the HeLa cell line. Henrietta Lacks, and the HeLa cell line that was established from her tumor cells, have made significant contributions to scientific progress and advances in human health. The authors declare no competing financial interest.

## Author contributions

S.M. Camus: conceptualization, investigation, formal analysis, supervision, writing-original draft, review and editing. M.D. Camus, C. Figueras-Novoa, G. Boncompain, L.A. Sadacca, D. Kioumourtzoglou: investigation, formal analysis. C. Esk, A. Bigot: resources. G.W. Gould, N.J. Bryant: conceptualization, supervision, writing-review and editing. F. Perez: conceptualization, supervision. S. Mukherjee: conceptualization, investigation, formal analysis, supervision, writing-review and editing. F.M. Brodsky: Conceptualization, supervision, formal analysis, funding acquisition, project administration, writing-original draft, review and editing.

## SUPPLEMENTARY INFORMATION

**Figure S1:**
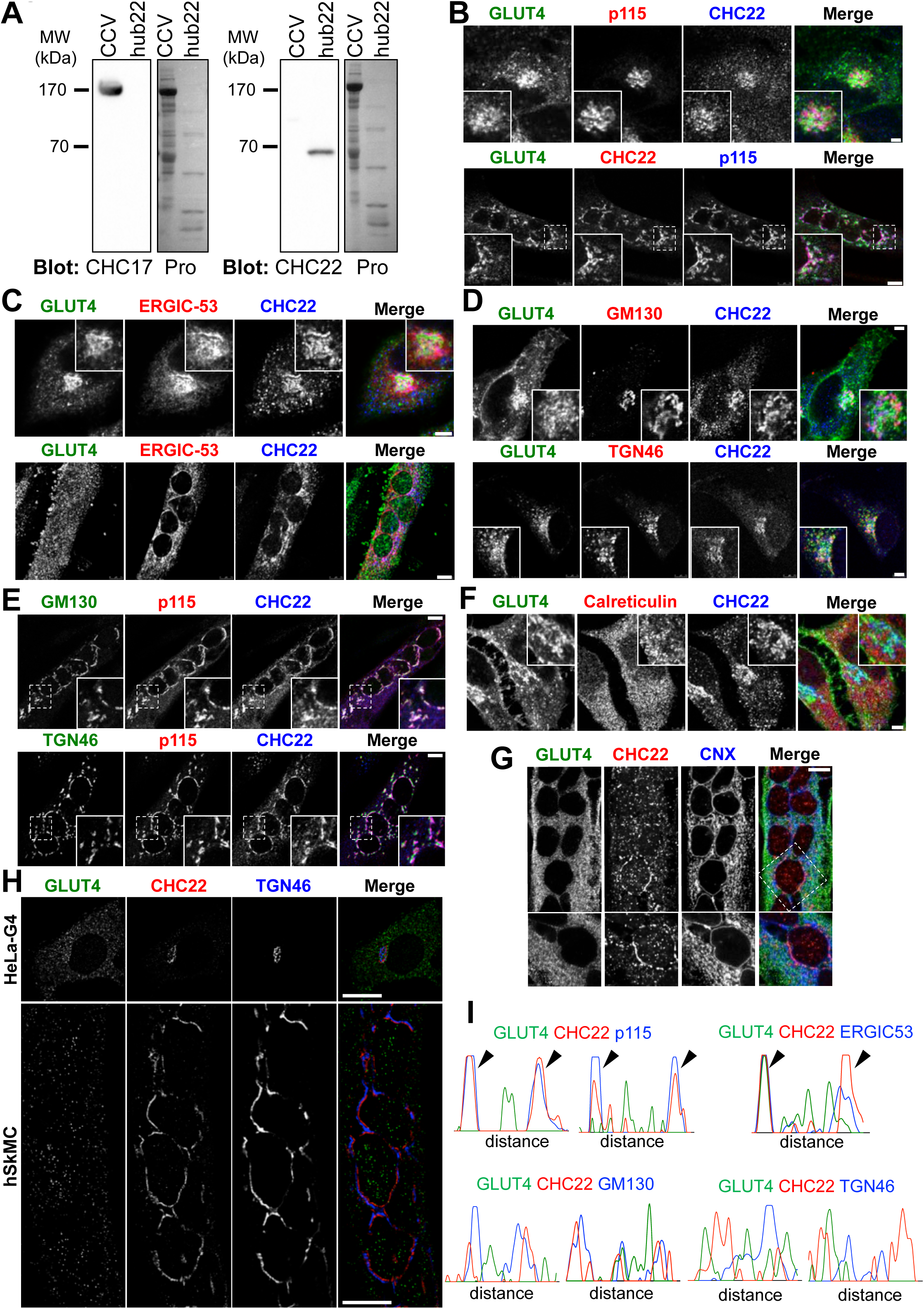
Immunofluorescence localization of CHC22 at the ER-to-Golgi Intermediate Compartment in HeLa-GLUT4 cells and in human skeletal muscle cells. (A) CHC17 (X22 antibody) or CHC22 (CLTCL1 antibody from Proteintech) immunoblots (IB) of clathrin coated vesicles (CCV) purified from pig brain containing only CHC17 or of cell lysate from bacteria expressing low levels of the hub fragment (residues 1074-1640) of CHC22 (hub 22). The migration of molecular weight (MW) markers is indicated in kilodaltons (kDa). Ponceau staining for proteins is shown on the right (Pro). (B) Representative confocal microscopy immunofluorescence (IF) imaging of CHC22 (red or blue), p115 (red or blue) and GLUT4 (green) in HeLa-GLUT4 cells (top panel) or LHCNM2 myotubes (bottom panel). (C) Representative IF staining for CHC22 (blue), ERGIC-53 (red) and GLUT4 (green) in HeLa-GLUT4 cells (top panel) or LHCNM2 myotubes (bottom panel). Scale bars: 5 µm for HeLa-GLUT4 cells and 7.5 µm for LHCNM2 myotubes in (B) and (C). (D) Representative IF staining for CHC22 (blue), GM130 or TGN46 (red) and GLUT4 (GFP, green) in HeLa-GLUT4 cells. Scale bars: 5 µm. (E) Representative IF staining for CHC22 (blue), GM130 or TGN46 (green) and p115 (red) in LHCNM2 myotubes. Scale bars: 7.5 µm. (F) Representative IF staining for CHC22 (blue), calreticulin (red) and GLUT4 (green) in HeLa-GLUT4 cells. Scale bars: 5 µm. (G) Representative IF staining for CHC22 (red), calnexin (CNX, blue) and GLUT4 (green) in hSkMC-AB1190-GLUT4. Scale bars: 10 µm. (H) Representative Structured Illumination Microscopy (SIM) of a HeLa-GLUT4 (HeLa-G4) cell (top panel) and human skeletal muscle cell (hSkMC-AB1190-GLUT4, bottom panel) stained for CHC22 (red) and TGN46 (blue). GLUT4 (green) was detected by GFP tag in HeLa-GLUT4 and immunostained with an anti-GFP antibody in hSkMC-AB1190-GLUT4. Scale bar: 10 µm. Merged images in (B-H) show red/green overlap in yellow, red/blue overlap in magenta, green/blue overlap in turquoise, and red/green/blue overlap in white. (I) Representative fluorescence intensity plots for GLUT4 (green), CHC22 (red) and p115, ERGIC-53, GM130 or TGN46 (blue) generated from SIM images of the perinuclear region of HeLa-GLUT4 cells.

**Figure S2:**
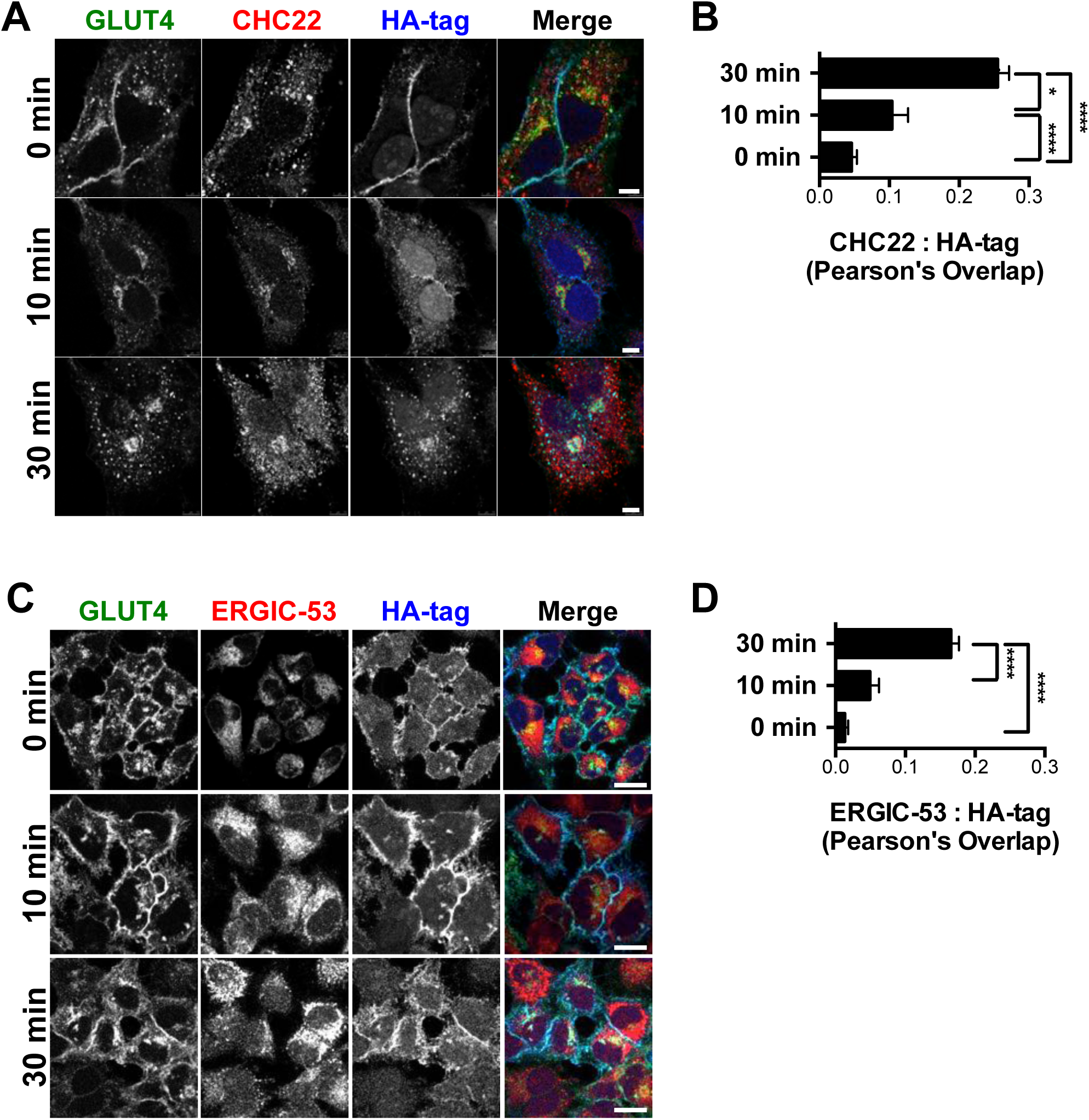
Surface GLUT4 is recycled to the GSC in proximity to the ERGIC. (A) Representative immunofluorescence (IF) staining for internalized surface-labeled GLUT4 (HA-tag, blue) and CHC22 (red) for HeLa-GLUT4 cells at 0, 10 or 30 minutes after insulin treatment. Total GLUT4 is detected by GFP tag (green). Scale bars: 5 µm. (B) Pearson’s overlap for labeling of CHC22 and HA-tag (data expressed as mean ± SEM, N=3, 8-40 cells per experiment). One-way analysis of variance (ANOVA) followed by Bonferroni’s multiple comparison post-hoc test *p<0.05, ****p<0.0001. (C) Representative immunofluorescence (IF) staining for internalized surface-labeled GLUT4 (HA-tag, blue) and ERGIC-53 (red) for HeLa-GLUT4 cells at 0, 10 or 30 minutes after insulin treatment. Total GLUT4 is detected by GFP tag (green). Scale bars: 10 µm. (D) Pearson’s overlap for labeling of ERGIC-53 and HA-tag (data expressed as mean ± SEM, N=3, 14-22 cells per experiment). One-way analysis of variance (ANOVA) followed by Bonferroni’s multiple comparison post-hoc test ****p<0.0001. Merged images show red/green overlap in yellow, red/blue overlap in magenta, green/blue overlap in turquoise, and red/green/blue overlap in white.

**Figure S3:**
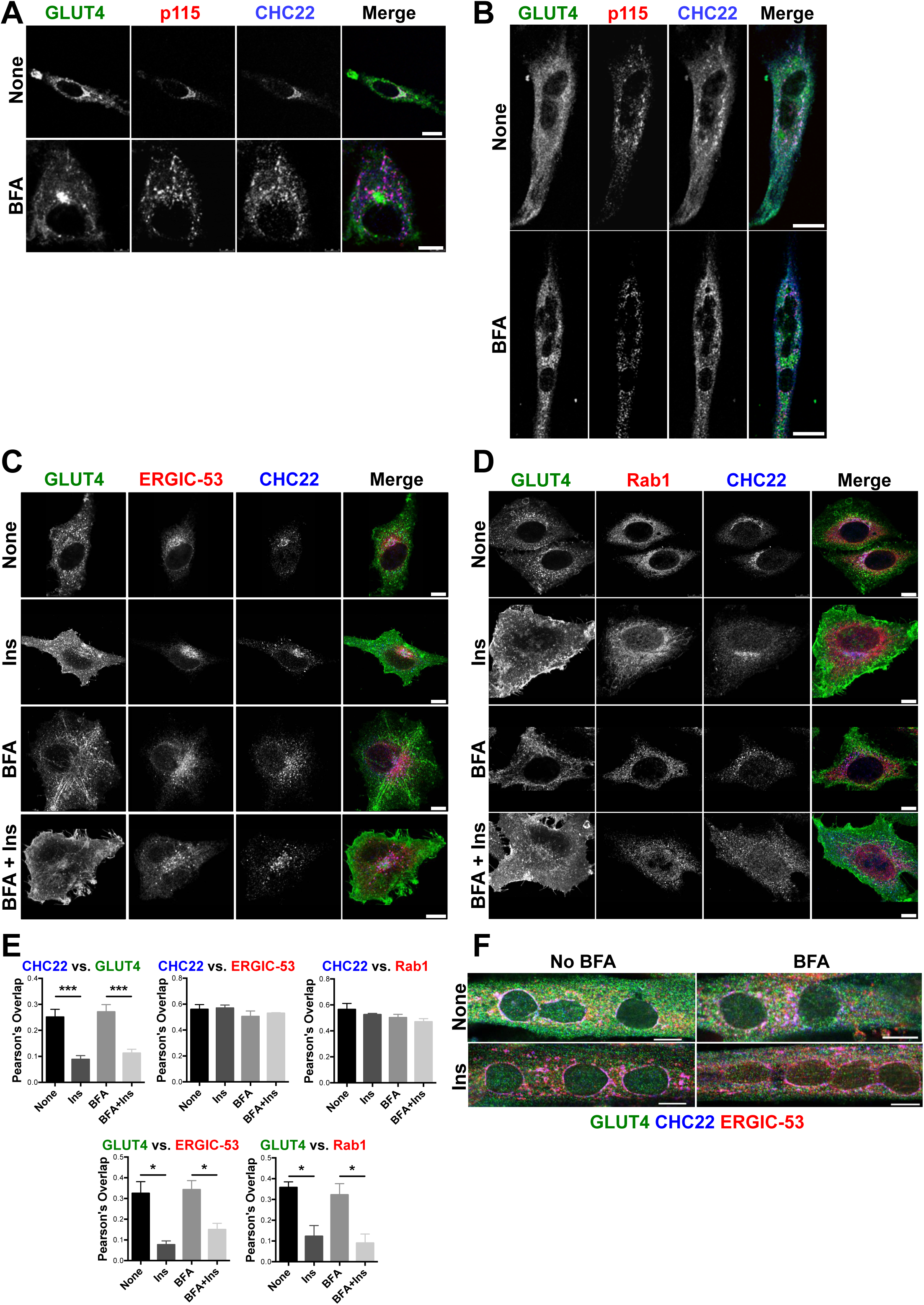
CHC22 re-distributes with p115 following Brefeldin A treatment. (A, B) Representative immunofluorescence (IF) staining for CHC22 (blue) and p115 (red) in HeLa-GLUT4 cells (A) or LHCNM2 myotubes (B) treated or not with Brefeldin A (BFA). GLUT4 (green) was detected by GFP tag in HeLa-G4 cells and by IF of endogenous protein in LHCNM2 cells. Scale bars: 5 and 25 µm for (A) and (B), respectively. (C, D) Representative immunofluorescence (IF) staining of HeLa-GLUT4 cells for CHC22 (blue) and ERGIC-53 (red) in (C) or Rab1 (red) in (D) treated or not with BFA and stimulated or not by insulin (Ins). GLUT4 (green) was detected by GFP. Scale bars: 10 µm. (E) Quantification of Pearson’s overlap values between CHC22, GLUT4, ERGIC-53 and Rab1 (data as in (C & D) expressed as mean ± SEM, N=3 to 4, 5 to 42 cells per experiment). One-way analysis of variance (ANOVA) followed by Tukey’s multiple comparison post-hoc test *p<0.05, ***p<0.001. (F) Representative immunofluorescence staining for CHC22 (blue), ERGIC-53 (red) and GLUT4 (anti-GFP antibody, green) in hSkMC-AB1190-GLUT4 treated or not with Brefeldin A (BFA) and stimulated or not by insulin (Ins). Scale bar: 10 µm. Merged images show red/green overlap in yellow, red/blue overlap in magenta, green/blue overlap in turquoise, and red/green/blue overlap in white.

**Figure S4:**
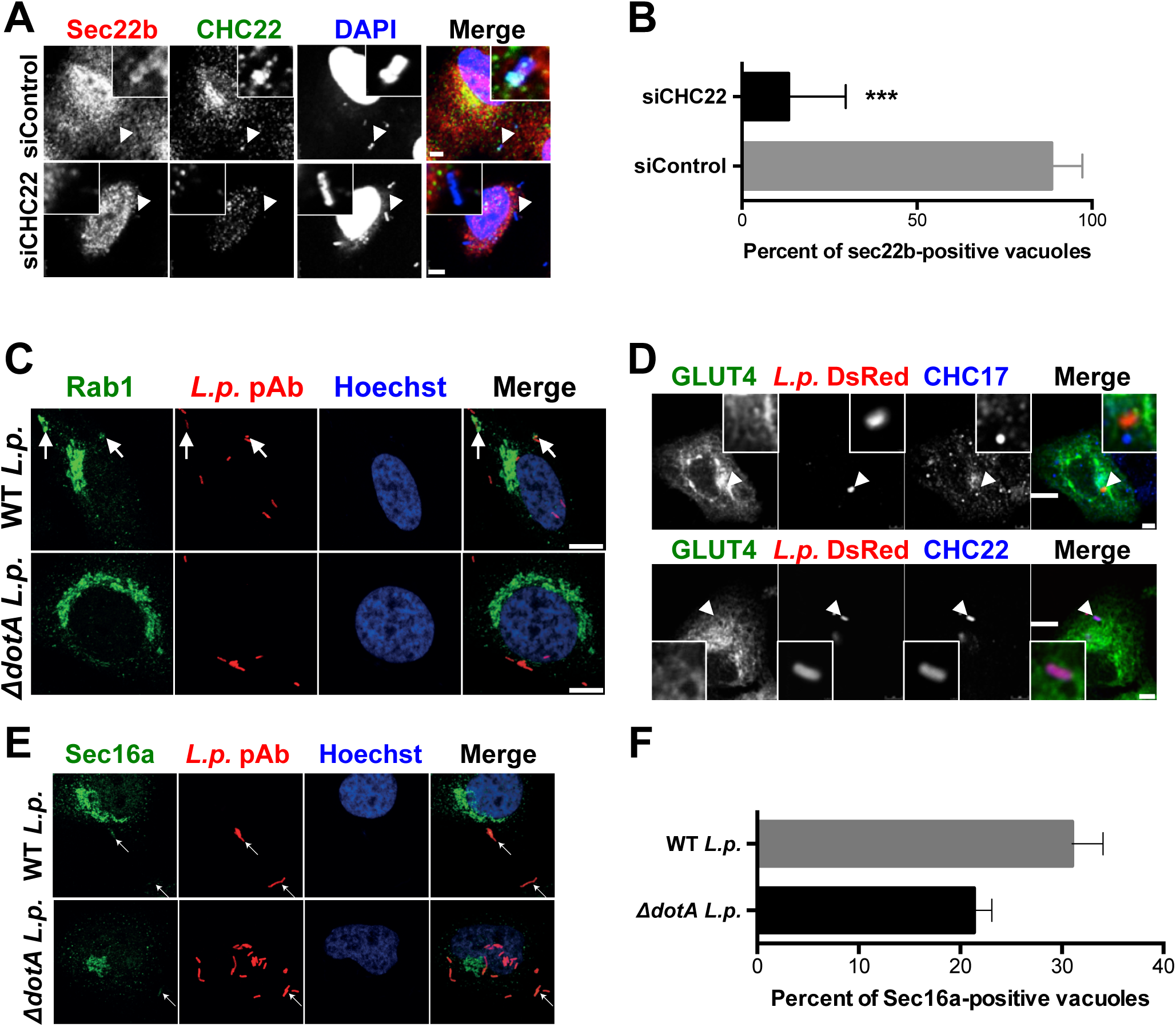
GLUT4 is not recruited to *Legionella pneumophila*’s replicative vacuole. Representative immunofluorescence images of single A549 cells from cultures treated with siRNA targeting CHC22 or with non-targeting control siRNA and labeled for Sec22b (red) and CHC22 (green), 1h post-infection with wild-type *L.p.* (MOI=50). Arrows point to *L.p.* detected with DAPI. Boxed inserts show *L.p.* region at 5X and 2X magnification for siControl and siCHC22, respectively. Scale bar: 5 µm referring to main images. (B) Quantification of the proportion of *L.p.* vacuoles staining positive for Sec22b. Data expressed as mean ± SEM, N=4, 10 to 20 vacuoles counted per experiment as represented in (A). Two-tailed unpaired Student’s t-test with equal variances ***p<0.001. (C) Representative images of HeLa cells transiently expressing FcγRII (needed for *L.p.* infection) (Arasaki and Roy, 2010), infected with wild type (WT) or mutant *ΔdotA L.p.* (MOI=50, red) and labeled 1 hour post-infection with antibodies against Rab1 (green). Hoechst stains the nuclei (blue). Arrows point to *L.p.*, Scale bar: 10 µm. (D) Representative images of A549 cells transiently transfected with HA-GLUT4-GFP (green), infected with wild type *L.p.* expressing mono-DsRed protein (*L.p.-*DsRed, MOI=50, red) and labeled 1 hour post-infection with antibodies against endogenous CHC17 (upper panel) or CHC22 (lower panel) (blue). Scale bars: 5 µm. (E) Representative images of HeLa cells transiently expressing FcγRII, infected with wild type (WT) or mutant *ΔdotA L.p.* (MOI=50, red) and labeled 1 hour post-infection with antibodies against Sec16a (E) (green). Hoechst stains the nuclei (blue). Arrows point to *L.p.*, Scale bar: 10 µm. (F) Quantification of the proportion of vacuoles staining positive for Sec16a. Data expressed as mean ± SEM, N=3, 50 vacuoles counted per experiment as represented in (E). Merged images in (A and D) show red/green overlap in yellow, red/blue overlap in magenta, green/blue overlap in turquoise, and red/green/blue overlap in white. Merged images in (C and E) show red/green overlap in yellow.

**Figure S5:**
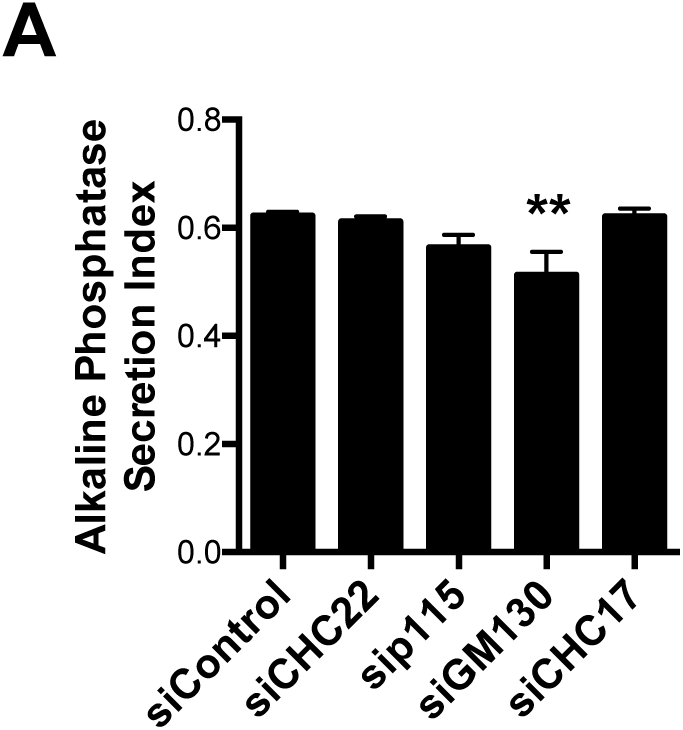
GM130 depletion affects the secretion of alkaline phosphatase in HeLa cells. Quantification of alkaline phosphatase secretion index for HeLa-GLUT4 cells treated with siRNA targeting CHC22, CHC17, p115 or GM130 or with non-targeting control siRNA. The alkaline phosphatase secretion index is the ratio of secreted enzyme activity (culture medium) to total cellular activity (secreted plus cell lysate). Data expressed as mean ± SEM, N=13-19 independent samples across 2 independent assays. One-way analysis of variance (ANOVA) followed by Bonferroni’s multiple comparison post-hoc test **p<0.01 versus siControl.

**Videos 1 and 2: Concurrent live video microscopy of GLUT1 and GLUT4 in HeLa cells indicates distinct trafficking pathways.**

Each video shows a HeLa cell expressing the endoplasmic reticulum (ER) Ii-hook fused to streptavidin along with HA-GLUT1-SBP-mCherry (GLUT1, red) and HA-GLUT4-SBP-GFP (GLUT4, green). The intracellular traffic of GLUT1-mCherry and GLUT4-GFP was simultaneously tracked for 1h after biotin addition released them from the ER. Upon ER exit, both GLUT1 and GLUT4 accumulated in the perinuclear region of the cell (yellow), then highly mobile GLUT1 vesicles become visible (red) while GLUT4 remained perinuclear. Timelapse videomicroscopy acquired on a spinning disk microscope at 8 frames per second. Video 1 stills are shown in Fig. 2A.

